# Reproductive history differentially shapes the neural response of middle-aged females to estradiol therapy after a metabolic challenge

**DOI:** 10.64898/2026.01.15.699773

**Authors:** Jennifer E Richard, Ahmad Mohammad, Stephanie E Lieblich, Kimberly A Go, Soda Wang, Rebecca K Rechlin, Tallinn FL Splinter, George E. Barreto, Liisa AM Galea

**Author notes:** Address all correspondence to: Liisa Galea, PhD, Treliving Family Chair in Women’s Mental Health, Senior Scientist, Centre for Addiction and Mental Health, Professor of Psychiatry, University of Toronto. These authors contributed equally (Jennifer E Richard, Ahmad Mohammad).

## Abstract

**Background:** Advancing age, the APOE□4 allele, and female sex are the top nonmodifiable risk factors for Alzheimer’s disease (AD). Female-specific experiences, such as parity and hormone therapy (HT) affect aging biomarkers such as metabolism and immune signaling and may even affect AD risk. Estradiol (E2), a common component of many HTs, affects cognition and brain health in aging females although this may vary depending on parity, genotype, and metabolic status. We hypothesized that prior parity influences brain and metabolic health, including response to E2, depending on APOE genotype.

**Methods:** Middle-aged female (10 month) wildtype (WT) or humanized (h) APOE□4 expressing rats, with different reproductive experience (nulliparous or primiparous) were fed a Western (WD) or standard diet (SD) for 2 months. In the second month, rats were given E2 or vehicle (oil) injections daily. Fear associative learning, plasma metabolic hormones, hippocampal inflammatory signalling, and neuroplasticity (neurogenesis, synaptic protein) were assessed.

**Results:** Females fed a WD gained weight and displayed metabolic dysregulation, regardless of genotype. E2 treatment reduced WD-induced weight gain and reduced metabolic hormones, with stronger effects in WT rats. E2 treatment increased dorsal hippocampal inflammatory signalling selectively in primiparous hAPOE□4 females fed a WD. Previous parity increased neurogenesis and reduced certain cytokines in the hippocampus of middle-aged WT rats under a SD. Both E2 treatment and previous parity decreased dorsal neurogenesis in hippocampus of hAPOE□4 rats. In hAPOE□4 females, higher weight was associated with reduced contextual fear memory, an effect driven by primiparous females. In the cued fear conditioning task, hAPOE□4 females displayed better cued fear memory than WT, however, WD exposure reduced cued fear memory in this group. Together, this indicates that diet and weight gain may be more detrimental to associative memory in hAPOE□4 females and that E2 treatment has more favourable outcomes in WT rats.

**Conclusions:** Previous parity alters how females respond to E2 and metabolic stress in midlife. Primiparous hAPOE□4 females were especially vulnerable to the effects of WD and E2, exhibiting more inflammation, impaired memory, and reduced weight-loss. These findings highlight the importance of considering parity and genotype when evaluating midlife metabolic and cognitive risk.

**Highlights:**

- Estradiol (E2) treatment reduced body weight gain under a Western diet (WD), with the most pronounced effects in wildtype primiparous rats.
- WD increased several metabolic hormones, and E2 treatment reduced several metabolic hormones after a WD in middle age, only in WT rats
- Increased body weight impaired contextual associative memory in primiparous hAPOE□4 females.
- E2 treatment increased dorsal hippocampal inflammatory signalling in primiparous hAPOE□4 rats and the WD increased inflammatory signalling in primiparous WT rats.
- Previous parity, but not E2 treatment, increased neurogenesis in WT rats only under standard diet conditions whereas a WD decreased neurogenesis based on genotype and previous parity

**Plain English Summary:** Alzheimer’s disease (AD) causes general cognitive decline, and females are at higher risk than males, particularly those carrying the APOEε4 gene. Female-specific experiences, such as previous pregnancy (parity) and hormone therapy, as well as lifestyle factors like body weight and diet, may further influence AD risk.

We examined how estradiol (E2), a hormone involved in the menstrual cycle, pregnancy, and some hormone therapies can affect memory, brain inflammation, synaptic plasticity (brain cell connectivity), and growth of new neurons (neurogenesis) in middle-aged female rats. We compared females with or without the APOEε4 gene and with or without prior parity, fed either a high-fat, high-sugar Western diet (WD) or a standard diet (SD).

WD increased body weight and worsened metabolic health, but the strength of negative effects depended on genotype and pregnancy history. In females with the APOEε4 gene and prior parity, higher body weight was linked to poorer memory. hAPOEε4 females on SD showed better memory than non-carriers but this benefit disappeared on WD, suggesting WD and obesity are particularly detrimental for hAPOE□4 carriers with prior parity. WD-fed primiparous hAPOE□4 females treated with E2 also showed higher brain inflammation than other groups. Pregnancy history and E2 treatment were associated with more favourable outcomes in females without the genetic risk for AD and fed a standard diet.

These findings highlight the importance of a personalized approach to hormone therapy and brain health in aging females. hAPOE□4 females with prior parity were most sensitive to the effects of diet. Together, this suggests genetic background, reproductive history, and diet interact to affect memory, inflammation, and neurogenesis, as well as the effects of E2 on these factors.

## Background

Alzheimer’s disease (AD) is a neurodegenerative disorder characterized by gradual and irreversible cognitive decline affecting millions globally (1). Risk factors for AD include non-modifiable factors, such as advancing age, female sex, and the presence of APOEε4 alleles. Many of the modifiable factors, however, are closely linked to metabolic health, including but not limited to obesity, diabetes, cardiovascular disease, and physical inactivity (2–7).

The lifetime risk of developing AD is approximately twice as high in females compared to males (5,6). Females with AD also exhibit more severe neuropathology and accelerated cognitive decline than males when diagnosed (8–11), which may be further exacerbated by metabolic factors (12). Furthermore, females with the APOEε4 allele have a greater risk to develop AD in middle age (13) and show greater central inflammation and cognitive decline than males (10,14–18). Given the heightened risk of AD in females, particularly among APOEε4 carriers, we investigated the potential contributing factors that influence AD risk using female-specific life experiences, such as previous parity and hormone therapy (HT) in middle-age, each of which impact metabolic and inflammatory systems (4,19–26). A common feature across these life events is changes in the levels of estrogens, such as estradiol (E2), a hormone that is critical for regulating brain function, metabolism, and immune responses, all factors which are individually implicated in AD pathophysiology.

The hippocampus, a critical region for learning and memory, is among the first areas to exhibit neuronal loss, and also shows reduced neurogenesis, and increased inflammation with AD (27–33). HT containing E2, given during the menopausal transition, can improve hippocampal-dependent cognition in postmenopausal females (34). However, HT’s ability to promote cognition can depend on the APOE genotype (35). E2 treatment can be beneficial in non-APOEε4 female carriers, whereas it may be detrimental in APOEε4 expressing female rodents and humans, however the results are very mixed (36–45). These mixed findings suggest that additional factors may influence the neuroprotective effects of E2, prompting us to investigate how body weight and parity history simultaneously contribute to the mixed findings of HT’s effects by APOE□4 genotype (46,47). Although, to our knowledge, no studies have interrogated the direct contribution of body weight, E2, and APOE genotype on AD endophenotypes. Lower BMI is associated with less brain aging in APOEe4 females (46,48) and a reduced risk of AD with menopause (49,50). Furthermore, HT’s impacts are affected by previous parity as nulliparous (never pregnant or mothered) rats show fewer beneficial neuroplastic effects after HT, than multiparous (pregnant and mothered multiple times) rats (51). These findings suggest that the effects of HT on the brain are influenced by both APOE genotype, reproductive history and potentially body weight.

Previous parity alters the trajectory of hippocampal aging in middle age in both humans and rodents (52). In humans, parity is associated with less brain aging regardless of the number of children in middle age (49, 50). Previous parity is also associated with increased hippocampal synaptic proteins, neurogenesis, and hippocampus-dependent cognition in middle age (51,54–63). Paradoxically, however, previous parity is also associated with a greater risk of AD and an earlier age of AD onset (55–58,64–66). The direction and magnitude of these effects depend on parity experience, as having 1-2 children may be beneficial against AD whereas having more than 3 children is associated with an increased risk of developing AD (56–58). In addition, previous parity in middle-aged hAPOEε4 female rats reduces hippocampal neurogenesis, activation of new neurons in response to memory retrieval, and increases pro-inflammatory cytokines in the hippocampus (67). Parity is also associated with profound changes to metabolic health that extend far beyond pregnancy (68–70). Taken together, parity influences hippocampal aging and AD risk in both humans and rodents differently depending on APOE genotype.

E2 plays a critical role in body-weight regulation, metabolism, appetite, and responsiveness to metabolic hormones such as insulin, with reductions of E2 increasing the risk of developing obesity and type II diabetes (71,72). The depletion of estrogens in middle age, together with the metabolic changes that occur at this time, collectively further increases the risk of AD (73), an effect exacerbated by exposure to diets containing high levels of saturated fats and refined carbohydrates, often referred to as a Western diet (WD; 2–4(74)). WD consumption increases inflammation and insulin resistance (75–80) and may contribute to impaired cognitive function (81), particularly during middle age (82–84). HT, containing E2, may help counteract these effects (85–87), however, whether parity or genetic background (e.g., APOE genotype) interact to alter the effects of E2 on metabolic health is unknown, in addition to its effects on inflammation, hippocampal plasticity and memory.

The aim of this study was to examine if E2 could mitigate effects of a metabolic challenge (WD consumption) on associative memory, inflammation, and neuroplasticity in middle aged females that differed by parity experience and APOE genotype. We hypothesized that prior parity would have a particularly detrimental impact on cognitive health in hAPOEε4 females, and that E2 treatment would yield genotype-specific outcomes. We hypothesize that there would be exacerbated deficits in hAPOEε4 females while providing benefits in wild-type counterparts, with these effects being more pronounced in previously parous animals.

## 1. Methods

### 1.1. Animals

Age-matched nulliparous or primiparous female wildtype (WT; n=39) or humanized (h) APOE□4 (n=41) knock-in (HsdSage:SD-*ApoE^em1Sage^* rat, developed by SAGE Labs, Inc., Saint Louis, MO, USA) Sprague Dawley rats were used. WT rats were used as controls in this study as an hAPOEε3 rat model has yet to be developed (88–90). Rats were pair-housed, except while pregnant or with their litter, in a 12-h light/dark cycle and had *ad libitum* access to water and standard diet (SD; Picolab®, rodent diet 5053). All protocols were approved by the Institutional Animal Care Committee at the University of British Columbia and conformed to the guidelines set out by the Canadian Council on Animal Care.

### 1.2. Breeding

At three months of age, half of the females were bred and produced a single litter (primiparous), while the remaining females remained nulliparous. To create primiparous rats, two females and one male of the same genotype (either both wild-type or both hAPOE□4 carriers) were housed together overnight. Each morning, vaginal lavage was performed on the females, and the samples were examined for the presence of sperm. If sperm was detected, the female was considered pregnant, weighed, individually housed in a clean cage, and monitored weekly throughout gestation. Litters were culled at 23 days, after which the females were re-grouped and housed together. Nulliparous rats were housed in a separate room to avoid sound and olfactory cues from primiparous rats and single housed for the same amount of time that the primiparous rats were housed by themselves.

### 1.3. Diet

At 10 months of age, nulliparous and primiparous WT and hAPOE□4 females were divided into weight-matched groups and fed either SD (Picolab®, rodent diet 5053) or western diet (WD; Research Diets (RD) Western Diet, D12079B, Research Diets Inc., New Brunswick, NJ, USA) for the following 7 weeks (until tissue collection).

### 1.4. Drugs

At 11 months of age, females of both genotypes, parity and both diet groups, were randomly divided into two groups receiving either daily subcutaneous injections of 17β-estradiol (E2; 0.3µg; E8875-250MG, MilliporeSigma. US) or sesame oil (vehicle; S3547-1L, Sigma-Aldrich, MilliporeSigma, US) at a volume of 0.1 mL for 3 weeks. Injections are preferred over implants due to variability in hormone levels with implants (91–93). The dose was chosen to mimic diestrus levels in rats which increase contextual fear memory and cell proliferation in the dentate gyrus (94,95).

### 1.5. Experimental timeline

At 11 months of age (after 4 weeks of SD or WD diet exposure), females of each genotype, parity and diet group were divided into weight-matched groups and given daily subcutaneous injections of vehicle (sesame oil; n = 5) or E2 (n = 4-6) as described. The timeline is illustrated in **Figure 1**. Rats were weighed twice weekly during the diet only period and daily prior to drug injection. Vaginal samples were collected via lavage during the diet and treatment periods.

**Figure 1.**
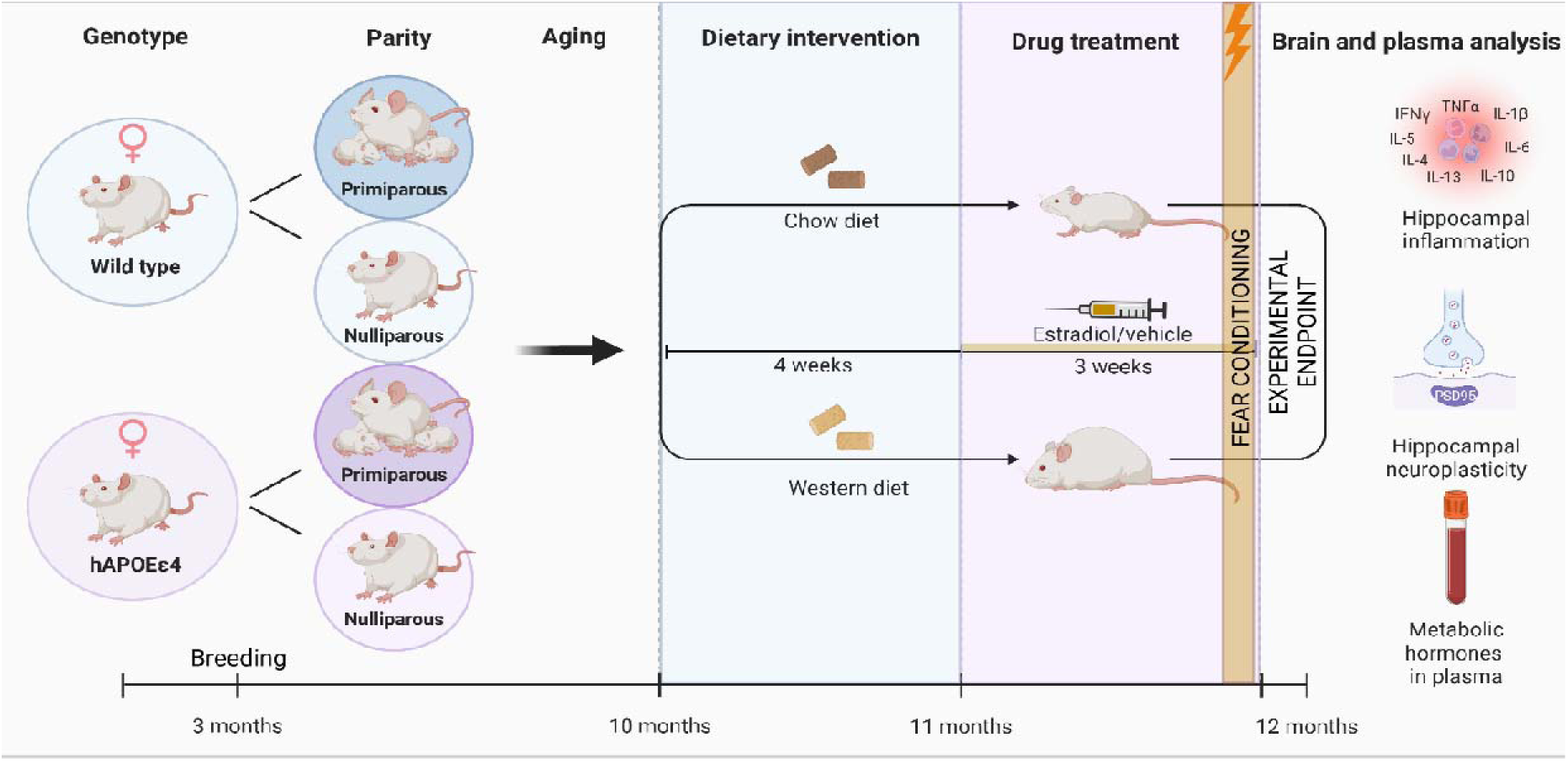
Graphical illustration of the experimental plan*. C*reated using BioRender.com.

### 1.6. Fear conditioning

After 3 weeks of hormone treatment (at approximately 12 months of age), rats underwent fear conditioning testing to assess hippocampus-dependent contextual fear conditioning and amygdala-dependent cued fear conditioning (96). The apparatus and protocol were as previously described (94). Briefly, conditioning and testing were performed in chambers (30.5 × 24 × 21□cm; Med Associates) equipped with a house light, speaker (4 kHz, 80□dB tone), and a grid floor connected to a shock generator (1.0□mA foot shocks). Behavior was recorded via overhead cameras, and background noise (70□dB) was maintained by ventilation fans. Locomotor activity was measured using an automated program (MED-PC, Med Associates), whereas freezing behavior was scored manually by experimenters blinded to the groups.

Training and testing occurred over two days. On day 1 (conditioning), rats received E2 or vehicle injections and, 30 minutes later, were transported in a clean cage by a different experimenter to Context A (bare aluminum and Plexiglass walls, lavender odor). After 3 minutes of exploration, rats received three tone–shock pairings (30 second tone co-terminating with a 2□second shock, 60□second inter shock intervals). Rats were returned to their home cage 1 minute after the final shock.

On day 2 (24□hours later), rats were again treated and transported to Context A for an 8-minute test of contextual fear (no tones or shocks). One hour later, they were brought via a novel route to Context B (striped wall inserts, tropical odor). After 3 minutes, three tones were presented (30 seconds each, 60 seconds apart) to assess cued fear memory. Freezing during each tone and the 3 minutes prior (context discrimination) was recorded.

### 1.7. Blood and brain collection

Food was removed from all cages one hour before tissue collection which occured during the rats’ light cycle (9:00AM-3:00PM). Trunk blood was collected via live decapitation and transferred into cold EDTA-coated tubes. Samples were centrifuged for 15 minutes, and plasma was stored at -80°C. Brains were rapidly extracted, and hemispheres were separated. The brain was flash-frozen on dry ice and stored at -80°C. Brain regions of interest (dorsal and ventral hippocampus) were micro-dissected from 50 µm sections using punchers in a cryostat maintained at -20°C. Cytokine levels were measured using electrochemiluminescence immunoassay kits as described below.

### 1.8. Hippocampal cytokine and PSD95 measurements

Electrochemiluminescence immunoassay kits (Meso Scale Discovery, Rockville, MD) were used to quantify cytokines (IFN-γ, IL-1β, IL-4, IL-5, IL-6, IL-10, IL-13, CXCL-1, and TNF-α; V-PLEX Proinflammatory Panel 2 Rat Kit, Cat# K15059D) and postsynaptic density protein 95 (PSD95; Cat# K150QND) in homogenized dorsal and ventral hippocampal samples. Tissue was homogenized using a TissueLyser III (Qiagen) with five beads in 200 µl of cold Tris lysis buffer (150 mM NaCl, 20 mM Tris, pH 7.4, 1 mM EDTA, 1 mM EGTA, 1% Triton X) supplemented with protease inhibitors (Complete Protease Inhibitor Cocktail, Roche), phosphatase inhibitors (Cocktails 2 and 3, Sigma-Aldrich), NaF, and PMSF in DMSO.

Homogenates were centrifuged at 100□rpm for 30□seconds at 4°C. Supernatants were aliquoted and stored at -80°C. Protein concentrations were determined using a BCA assay kit (Thermo Scientific). Samples were diluted 1:10 for PSD95 and 1:4 for cytokine assays in Tris lysis buffer and run in duplicate per manufacturer instructions. For cytokine plates, sample and calibrator incubation was extended to overnight at 4°C (vs. 2 hours). Plates were read using a MESO QuickPlex SQ 120 (Meso Scale Discovery), and data were analyzed with Discovery Workbench 4.0. Cytokine concentrations were calculated from standard curves and expressed as pg/mg protein; PSD95 values were normalized to protein and expressed as signal/pg□mg□¹.

### 1.9. Plasma hormone levels

Metabolic hormones (C-peptide, ghrelin, GLP-1, glucagon, insulin, leptin, and PYY) were measured in plasma using U-PLEX Metabolic Group 1 Rat Kits (Meso Scale Discovery, Rockville, MD; Cat# K15200L). Plasma samples were diluted 1:2 with the Metabolic Assay Working Solution and run in duplicate according to the manufacturer’s instructions. Plates were read using a MESO QuickPlex SQ 120 (Meso Scale Discovery), and data were analyzed with Discovery Workbench 4.0 software. Hormone concentrations were calculated from standard curves and reported as pg/ml.

### 1.10 Immunohistochemistry

A series of free-floating tissue was rinsed 3 times at room temperature with 0.1 M phosphate-buffered saline (PBS) for 10 minutes each. The tissue was then incubated in 10% Triton X for 30 minutes. Sections were added to the blocking solution (10% normal donkey serum (NDS) and 3% Triton X in PBS) for 1 hour. The tissue was immediately incubated in rabbit anti-doublecortin (DCX; 4604S, Cell Signal Technology Inc. Danvers) diluted 1:1000 in blocking solution (10% NDS and 3% Triton X in PBS) for ∼48 hours at 4°C. Following PBS rinses (10 minutes X3), tissue was incubated for 4 hours in Alexa Fluor 488 conjugated donkey anti-rabbit (1:500; A-21206, Thermo Fisher Scientific) in a solution PBS containing 10% NDS, 3% Triton-X. Sections were then washed in PBS for 3x 10 minutes and then stained with 4,6-diamidino-2-phenylindole (DAPI; 1:500; D1306, Thermo Fisher Scientific) for 2.5 minutes at room temperature. The tissue was rinsed for 5 minutes (3X). Finally, tissue was mounted onto Superfrost/Plus slides (Fisher scientific) and cover slipped using PVA-DABCO mounting medium.

### 1.11 DCX-expressing cells: quantification and microscopy

DCX-expressing cells were imaged with Zeiss Axio Scan.Z1 (Carl Zeiss Microscopy, Thornwood, NY, USA) with a 20x objective using fluorescent imaging. DCX-expressing cells were then quantified by an investigator blinded to experimental conditions. Two dorsal and two ventral sections were quantified for each rat. The density was calculated by dividing the number of DCX-expressing cells with the area of the dentate gyrus for each section.

Photomicrographs were acquired using the Fluoview 4000 confocal microscope (Evident Scientific) with a 20x objective lens. DAPI was visualized using the 405nm laser (blue) and DCX was visualized using the 488nm laser (green).

### 1.12 Statistical analyses

Data are presented as mean ± standard error of the mean (SEM). Statistical analyses were performed using TIBCO Statistica (v. 9, StatSoft, Inc., Tulsa, OK, USA) with analyses of variance (ANOVA), including genotype (WT, hAPOE□4), parity (nulliparous, primiparous) diet (SD, WD), and treatment (sesame oil, E2) as between-subject factors for each dependent variable. For body weight, weeks was added as the within-subjects factor. Graphs were generated using GraphPad Prism (GraphPad Software, Inc., San Diego, CA). Effect sizes are reported for significant results, including partial eta squared (ηp²) or Cohen’s d when appropriate. Post hoc comparisons were conducted using Newman-Keuls tests, and a priori comparisons used least significant difference (LSD) and were subjected to Bonferroni corrections. Principal component analysis (PCA) was conducted on cytokine profiles from hippocampal tissue and metabolic hormones from serum. Before running the principal component analyses (PCA), we standardized all variables to z□scores (mean = 0, SD = 1) to ensure comparability across measures. PCA was performed without rotation, and component scores were calculated using the regression method. ANOVA models were applied to the first two principal component scores to quantify variance explained by cytokine and metabolic profiles in circulating and neural tissue, as previously described (97). Linear regression analyses were conducted to examine relationships between body weight and freezing behavior during contextual and cued fear conditioning, with separate models run for each genotype and parity group to assess potential moderating effects. Statistical significance was set at p ≤ 0.05.

## 2. Results

### Western diet-induced weight gain is reduced with E2 treatment in middle-aged female rats with stronger effects in primiparous WT animals

As expected, WD consumption for 4 weeks led to a significant increase in body weight compared to SD regardless of genotype or previous parity status (main effect of Diet: F_(1, 73)_ = 160.603, *p* < 0.001,□p^2^ = 0.688; Weeks x Diet interaction: F_(3, 219)_ = 28.035, *p* < 0.0001,□p^2^ = 0.277. ; **Figure 2A**). There were no other significant main effects or interactions present (*p* ≥ 0.089).

**Figure 2.**
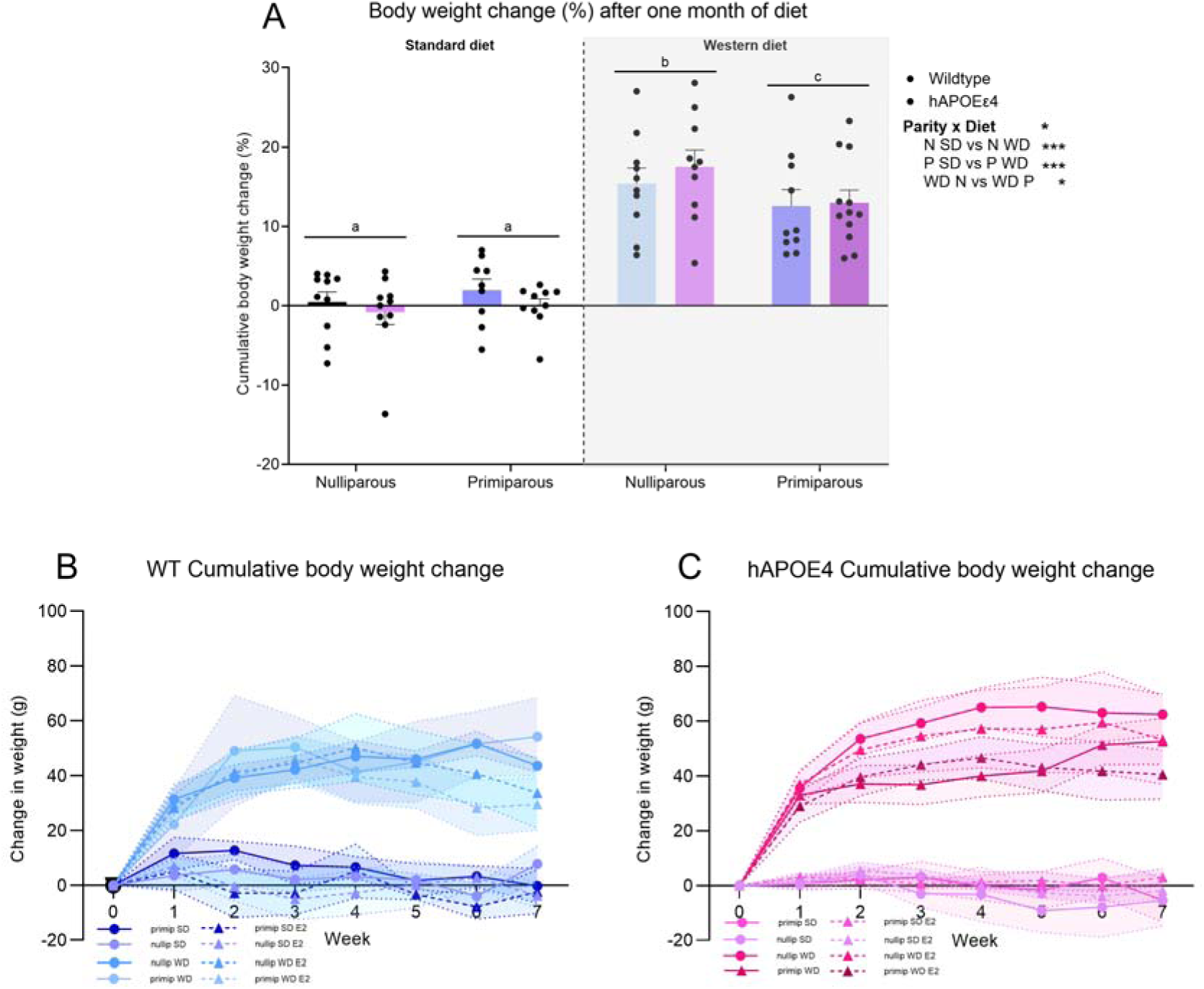
Western-diet (WD) consumption increases body weight and estradiol (E2) reduces body weight in WD females at middle age. Percent body weight on diet (A) total body weight change in grams is displayed in (B) for WT females and (C) for APOE□4 females. n = 9-10 females per genotype, parity and diet for diet only groups and 3-6 for treatment, genotype, parity and diet groups. * p < 0.05, ** p < 0.01, ***p < 0.001, ****p < 0.0001, # p < 0.10. N = nulliparous; P = primiparous; E2 = estradiol.

**Figure 2.**
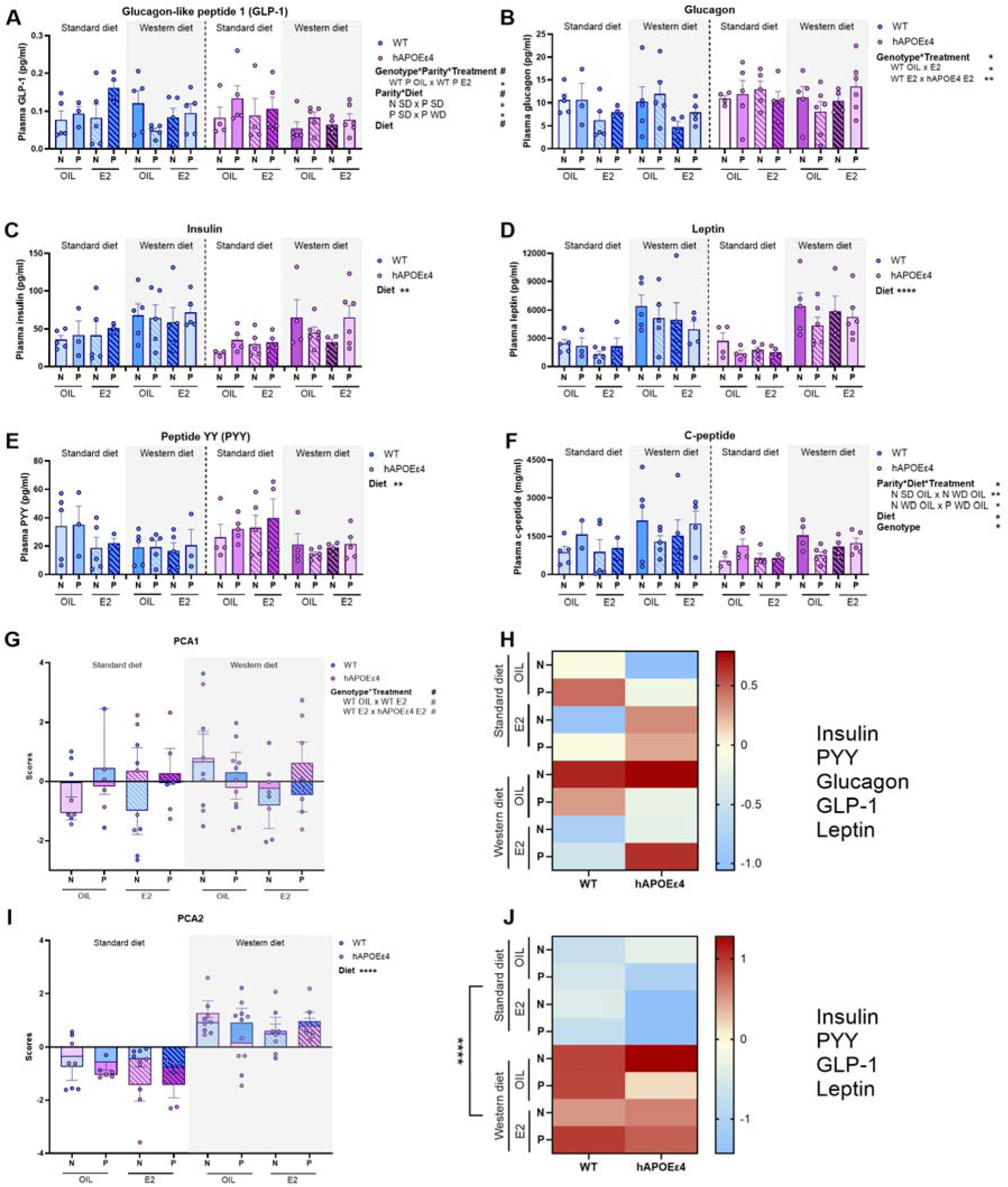
Western-diet (WD) increased the levels of insulin, leptin and C-peptide, as well as reducing peptide YY (PYY), in plasma. Estradiol (E2) treatment reduces plasma glucagon and humanized APOE□4 (hAPOE□4) genotype was associated with elevated levels of c-peptide compared to wildtype (WT). Plasma hormone levels of glucagon-like peptide-1 (GLP-1; A), glucagon (B), insulin (C), leptin (D), peptide YY (PYY; E), C-peptide (F). Principal component analysis (PCA) and heatmap for component 1 (G) and component 2 (H). Estrous cycling was similar between genotypes at 10-11 months of age (I). n = 9-10 females per genotype, parity and diet for diet only groups and 3-6 for treatment, genotype, parity and diet groups. * *p* < 0.05, ** *p* < 0.01, \*\*\**p* < 0.001, \*\*\*\**p* < 0.0001, # *p* < 0.10. R = regular cycling; IRR = Irregular cycling; ACY = Acyclic; N = nulliparous; P = primiparous; E2 = estradiol.

In the last 3 weeks of diet exposure and treatment, under WD fed conditions only, E2 reduced body weight gain, with stronger effects in primiparous rats. In primiparous rats, weight gain was reduced after two and three weeks of E2 treatment (both p’s<0.001; after one week of treatment p=0.439). In nulliparous rats, E2 reduced body weight gain only after 3 weeks of treatment (p=0.032, one and two weeks of E2 treatment-p=0.610, p=0.303, respectively). There were no significant effects in SD fed rats with E2 treatment (all p’s >0.676). Weeks by Treatment x Diet x Parity interaction: F_(1, 65)_ = 3.104, *p* = 0.048,□p^2^ = 0.045; **Figure 2B and C**). A priori, we were interested in genotype influences and the effects of E2 treatment to reduce body weight gain were only seen in the WT primiparous group (all ps <0.00001) but not the nulliparous WT or hAPOE□4 groups (**Figure 2B and C)**. There was also a main effect of diet (Treatment: F_(1, 65)_ = 130.49, *p* < 0.001,□p^2^ = 0.668), and a week by Treatment effect (F_(1, 130)_ = 3.428, *p* = 0.0354,□p^2^ = 0.050) but there were no other significant main effects or interactions (*p* ≥ 0.054).

### 2.1 Western diet consumption increased several metabolic hormones and E2 treatment reduced glucagon in WT but not in hAPOEe4 rats fed a WD

To assess the metabolic impact of 7 weeks of WD exposure, we measured the levels of several metabolic hormones in plasma. WD consumption led to increased levels of plasma insulin (Diet: F_(1, 58)_ = 8.692, *p* = 0.005, □p^2^ = 0.130; **Figure 3C**), leptin (Diet: F_(1, 60)_ = 29.631, *p* < 0.000, □p^2^ = 0.331; **Figure 3D**), and C-peptide (Diet: F_(1, 50)_ = 4.242, □p^2^ = 0.078, *p* = 0.045; **Figure 3F**) but reduced levels of peptide YY (PYY; Diet: F_(1, 56)_ = 9.765, *p* = 0.003, □p^2^ = 0.148; **Figure 3E**) and GLP-1 (trend: Diet: F_(1, 59)_ = 3.631, *p* = 0.062, □p^2^ = 0.0580; **Figure 3A**), regardless of past parity or genotype.

**Figure 3.**
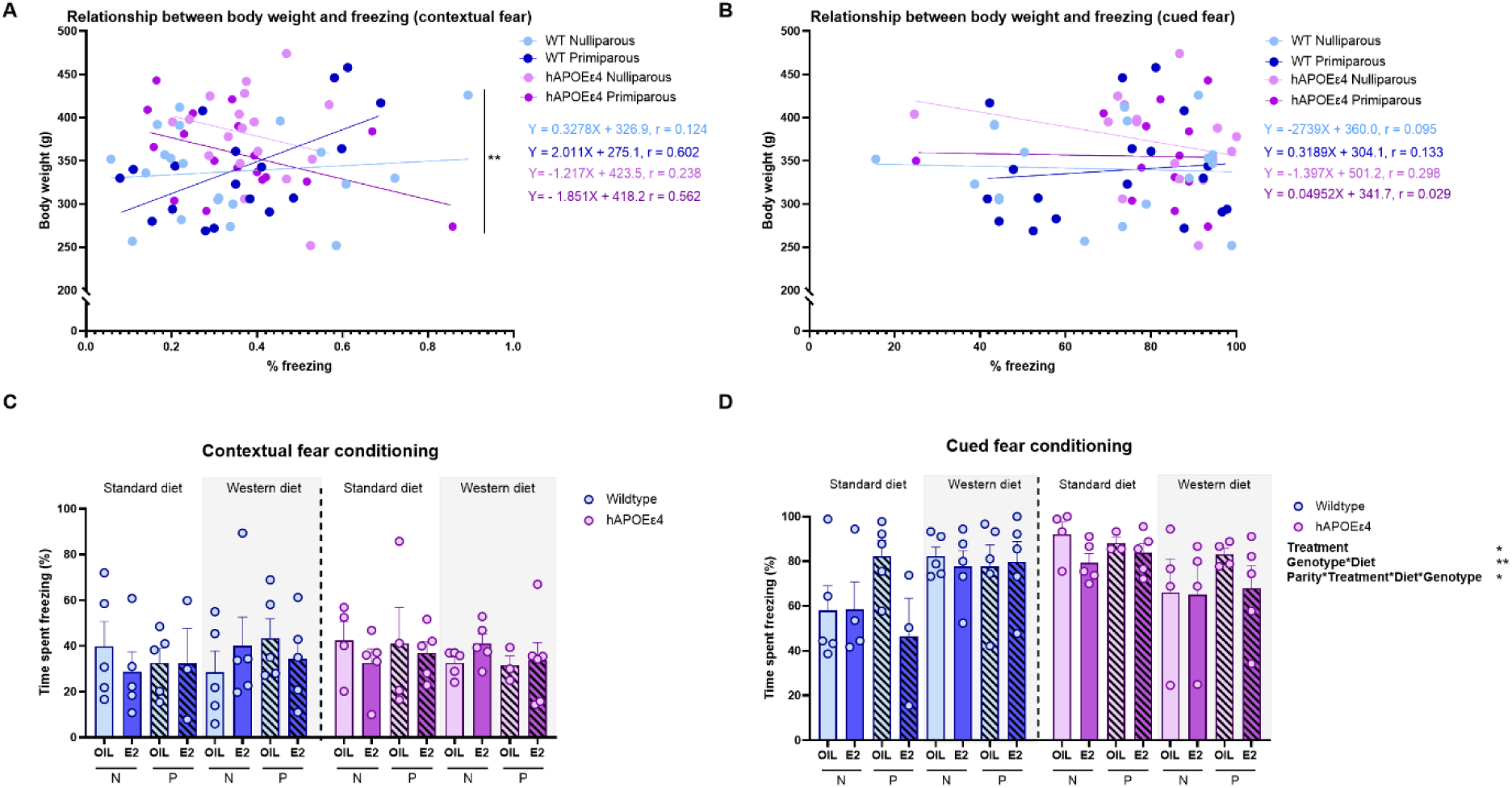
Linear regression shows a positive relationship between body weight and freezing in the contextual fear test in wild-type (WT) females at middle age, whereas a trend towards a negative relationship is seen in hAPOE□4 females; these effects are driven by females with previous parity. The relationship between body weight and freezing between genotypes with and without previous parity in the contextual fear conditioning (A) and cued fear conditioning task (B). Percent freezing in the contextual fear conditioning (C) and cued fear conditioning tasks (D). n = 9-12 females per genotype and parity and 3-6 for treatment, genotype, parity and diet groups. * p < 0.05, ** p < 0.01, # p < 0.10. E2 = estradiol, WT = wildtype, SD= standard diet, WD = western diet.

For C-peptide, hAPOE□4 females had lower levels than WT (Genotype: F_(1, 50)_ = 5.747, □p^2^ = 0.103, *p* = 0.020), and there was also a Parity x Diet x Treatment interaction: F_(1, 50)_ = 4.832, *p* = 0.033, □p^2^ = 0.088) but no significant post-hoc tests emerged. E2 treatment in WT females led to reduced levels of glucagon compared to oil-treated WT females (*p* = 0.028, Cohen’s *d* = 0.935) and to E2-treated hAPOE□4 females (*p* = 0.006, Cohen’s *d* = 1.419); Genotype x Treatment interaction: F_(1, 60)_ = 6.078, *p* = 0.017, □p^2^ = 0.092; **Figure 3B**) an effect that was not present in hAPOE□4 females (*p* = 0.533). There were no other significant main or interaction effects; there were no other significant main effects or interactions (*p* ≥ 0.068).

Lastly, we conducted a PCA to see how diet, genotype, parity, and E2 treatment may impact these metabolic hormones collectively. The first two principal components (PC) explained 68.2% of the total variance in the dataset (PC1: 39.9%, PC2: 28.2%). Insulin, PYY, glucagon, GLP-1, and leptin all loaded positively onto PC1. In PC2, leptin and insulin loaded positively, whereas PYY and GLP-1 loaded negatively. An analyses of the loadings from the PCA of metabolic factors revealed a trend towards an interaction between genotype and treatment in which E2 treatment reduced the PC1 score in WT females (*p* = 0.066, Cohen’s *d* = 0.698), which was not seen in hAPOE□4 female (*p* = 0.403 ; Genotype x Treatment interaction: F_(1, 50)_ = 2.937, *p* = 0.093, □p^2^ = 0.055,; **Figure 3G**). Furthermore, there was a trend towards higher scores in PC1 of E2-treated hAPOE□4 females compared to E2-treated WT females (Cohen’s *d* = 0.711, *p* = 0.067). In PC2 we observed that WD resulted in increased scores compared to SD (Diet: F_(1, 50)_ = 47.638, □p^2^ = 0.488, *p* < 0.000; **Figure 3H**).

It was observed that 80% of females were irregular or acyclic during diet and treatment manipulations (**Figure 3I**) with similar proportions of cycling between wildtype and hAPOE□4 (χ² = 0.08, *p* = 0.78). The overall distribution of cycle types did not differ significantly between genotypes (χ² = 0.04, *p* = 0.98).

### 2.2 Weight gain and WD exposure are particularly detrimental to associative memory in parous hAPOE□4 females

There were no significant main or interaction effects for contextual fear conditioning (all *p* values >0.22); **Figure 4C**). In the cued fear conditioning task, post-hoc analyses revealed that in rats fed a WD compared to a SD increased freezing in WT females (*p* = 0.041, Cohen’s *d* =0.783) but decreased freezing in hAPOE□4 females (trend only *p* = 0.065 Cohen’s *d* = 0.759; Diet x Genotype interaction: F_(1, 55)_ = 12.974, *p* = 0.0007, □p² = 0.191; **Figure 4D**). Furthermore, on a SD hAPOE□4 females displayed more freezing than WT (*p* = 0.0037, Cohen’s *d* = 0.836), however, this effect was absent after WD exposure (p>0.174, Cohen’s *d* =0.627). There were trends for a main effects of treatment (F_(1, 55)_ = 3.651, *p* = 0.061) and Genotype (F_(1, 55)_ = 2.932, *p* = 0.092) and for a four-way interaction (F_(1, 55)_ = 3.135, *p* = 0.082; **Figure 4D**) but no other significant main effects or interactions were present (*p* ≥ 0.343).

**Figure 4.**
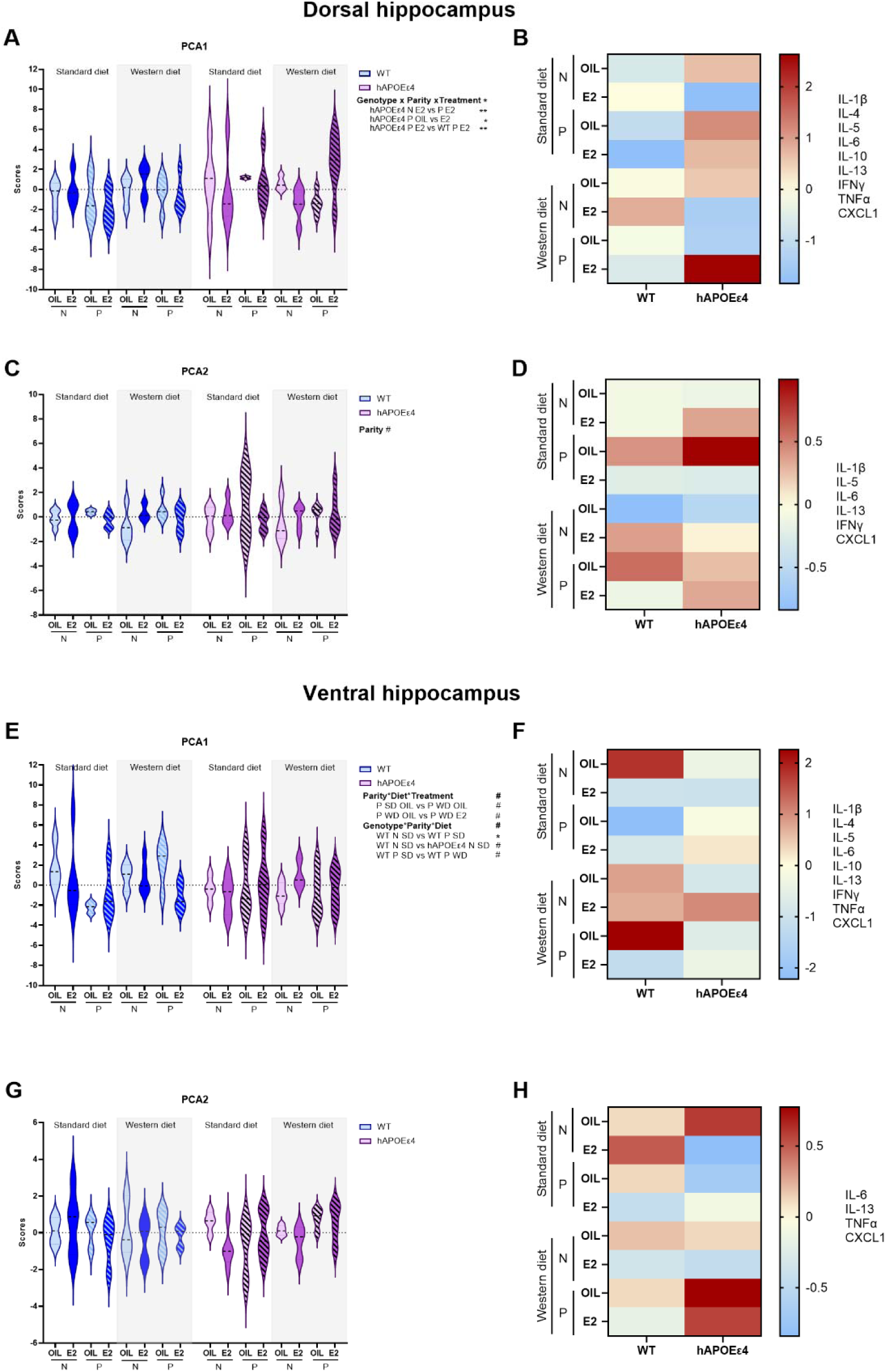
Principal Component Analysis (PCA) of inflammation in the dorsal and ventral hippocampus. Principal Component (PC)1 (A) and PC2 (C) in the dorsal hippocampus and a heat map for PC1 (B) and PC2 (D). PC1 (E) and PC2 (G) in the ventral hippocampus and a heat map for PC1 (F) and PC2 (H). n = 3-6 for treatment, genotype, parity and diet groups. *\** p *< 0.05, \*\** p *<* 0.01, # p < 0.10. E2 = estradiol, WT = wildtype, SD= standard diet, WD = western diet, N = nulliparous, P = primiparous.

Next, we aimed to determine whether body weight affects associative memory. Linear regression analyses were performed to examine the relationship between body weight and freezing behavior during contextual and cued fear conditioning across all groups. Importantly, body weight did not impact locomotor activity during the tasks (**Supplementary Figure 5**). To assess whether the association between body weight and contextual fear conditioning is modulated by genotype and reproductive history, linear regression analyses were conducted separately for WT and hAPOE□4 female rats, grouped by parity. In both nulliparous WT and hAPOE□4 females, no significant relationships were found between body weight and freezing in the contextual fear conditioning task (n = 20, R² = 0.013, F_(1,18)_ = 0.242, *p* = 0.629; hAPOE□4 n = 18, R² = 0.052, F_(1,16)_ = 0.858, *p* = 0.475; **Figure 4A**). However, in primiparous females, body weight and freezing behaviour relationships differed by genotype. In WT primiparous rats, body weight significantly predicted freezing behavior (n = 19, R² = 0.330, F_(1,16)_ = 7.867, *p* = 0.013), indicating that higher body weight was associated with increased contextual fear memory.

However, in hAPOE□4 primiparous females, body weight significantly predicted lower freezing behavior or reduced contextual memory (n = 18, R² = 0.224, F_(1,16)_ = 4.607, *p* = 0.048). A comparison of the regression slopes across groups revealed a significant difference (F_(3,66)_ = 4.032, *p* = 0.011), indicating that the relationship between body weight and contextual freezing behavior significantly varied depending on genotype and parity. Together, these findings suggest that body weight is differentially associated with contextual fear memory in WT and hAPOE□4 females, with opposing effects in primiparous animals.

Next, we performed linear regressions to assess the relationship between body weight and cued freezing behavior. Analyses stratified by genotype and parity (nulliparous vs. primiparous) similarly revealed no significant associations between body weight and cued freezing in any group: WT nulliparous (n = 18, R² = 0.001, F_(1,17)_ = 0.0178, *p* = 0.895), WT primiparous (n = 17, R² = 0.072, F_(1,17)_ = 0.017, *p* = 0.355), hAPOE□4 nulliparous (n = 17, R² = 0.090, F_(1,15)_ = 1.277, *p* = 0.277), or hAPOE□4 primiparous (n = 16, R² < 0.001, F_(1,14)_ = 0.008, *p* = 0.929; **Figure 4B**). Comparison of regression slopes across these four groups indicated no significant difference (F_(3,57)_ = 0.427, *p* = 0.659), and no significant differences were observed between intercepts (F_(3,60)_ = 1.384, *p* = 0.308).

Collectively, these data indicate that, at middle age, previous parity influences contextual fear memory differentially by genotype and body weight, with enhanced memory with increased body weight in WT, but reduced memory with hAPOE□4 genotype. In addition, hAPOE□4 females can perform better on cued memory compared to WT females, however this effect is lost with exposure to WD indicating that hAPOE□4 females may be more sensitive to metabolic changes at this time.

### 2.3 E2 increases dorsal hippocampal cytokine signalling in parous hAPOE□4 females, particularly under WD conditions

Given the complex and interrelated nature of inflammatory signaling, we performed PCAs on cytokine data from the dorsal and ventral hippocampus separately to reduce dimensionality and identify overarching patterns of inflammation signaling. Individual cytokine levels and analyses can be found in the Supplement. Because every cytokine, both pro□ and anti□inflammatory, loaded positively on PC1, we interpreted higher PC1 values as indicating a stronger overall cytokine activity or “inflammatory load,” rather than a shift toward a specific immune polarity. In the dorsal hippocampus, the first principal component (PC1) accounted for 55.6% of the variance and was strongly associated with elevated levels of multiple anti-inflammatory (IL-4, IL-10, IL-13) and pro-inflammatory (IL-1β, TNFα, IFNγ, IL-6) cytokines, suggesting it reflects a broad inflammatory signalling. PC2 explained an additional 13.0% (68.6% explained variance combined) of the variance and was most strongly correlated with IL-6, as well as showing distinct loadings for IL-5 and IL-13. Using the loadings, an ANOVA on the PCA found that E2 treatment in primiparous hAPOE□4 females led to higher scores in PC1 in the dorsal hippocampus than vehicle-treated primiparous hAPOE□4 females (*p* = 0.027, Cohen’s *d* = 1.057; Genotype x Parity x Treatment interaction: F_(1,55)_ = 4.674, *p* = 0.035□p^2^ = 0.078; **Figure 5A**) and all other groups nulliparous hAPOE□4 E2-treated females: *p* = 0.009, Cohen’s *d* = 0.927; primiparous WT E2-treated females: Cohen’s *d* = 1.207, *p* = 0.007). Most dramatic PCA increases were seen in WD-fed primiparous hAPOE□4 females receiving E2 as demonstrated in the heat map in **Figure 5B**. In PC2, we observed a trend towards increased scores in primiparous, compared to nulliparous, females (Parity: F_(1,55)_ = 3.576, *p* = 0.064; □p^2^ = 0.061; **Figure 5C** & **D**). There were no other significant interactions or main effects (*p* ≥ 0.098). Statistical details for PCA loadings for individual cytokines can be found in the Supplementary table 1.

**Figure 5.**
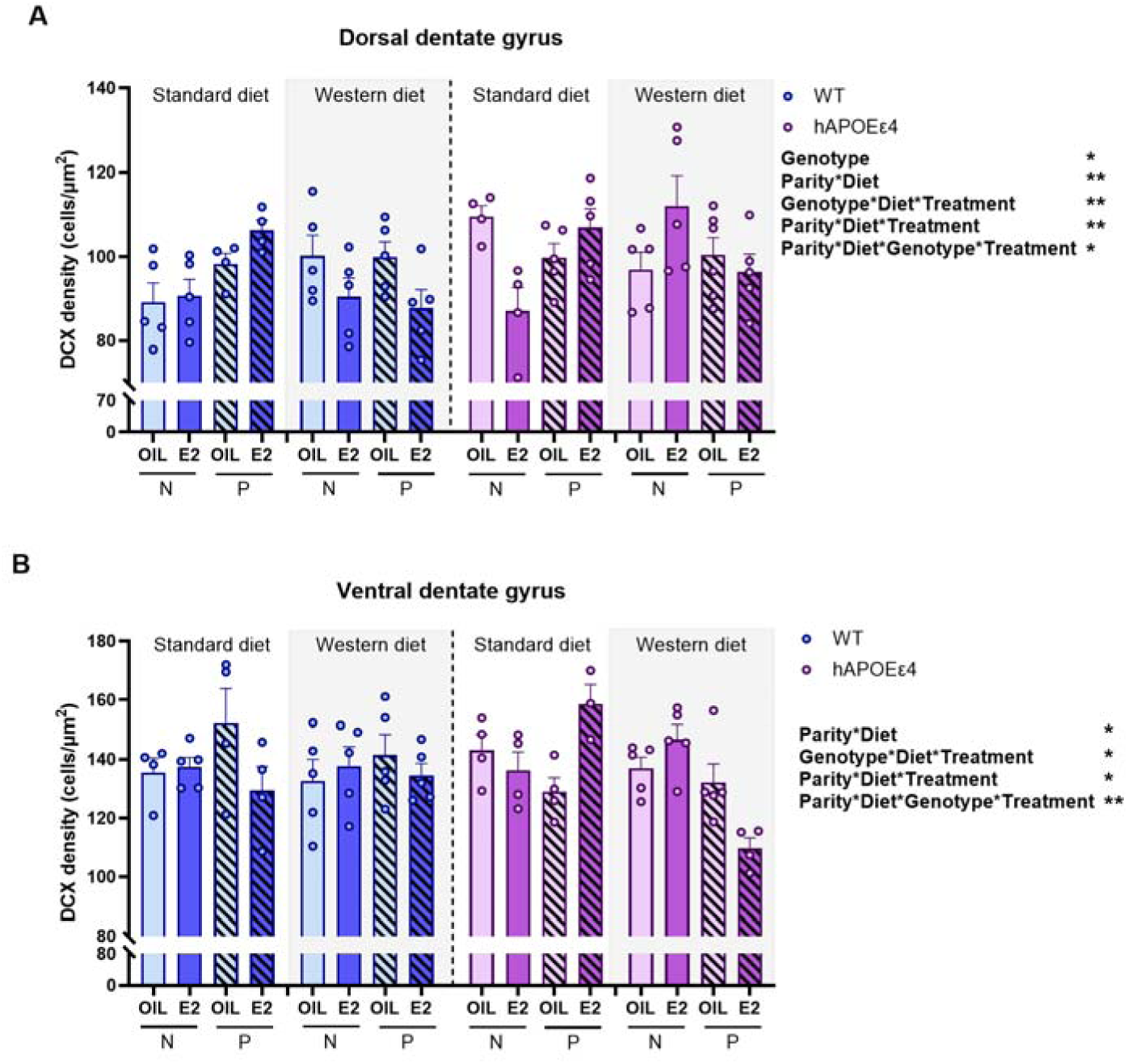
Effects of genotype, parity, diet, and E2 treatment on neurogenesis in the hippocampus. DCX density in the dorsal (A) and ventral (B) dentate gyrus in middle-aged females. n = 4-6 for treatment, genotype, parity and diet groups. * p < 0.05, ** p < 0.005 # < 0.100. E2 = estradiol, WT = wildtype, SD= standard diet, WD = western diet, N = nulliparous, P = primiparous.

In the ventral hippocampus, PC1 accounted for 56.8% of the variance and was strongly correlated with both anti-inflammatory cytokines (IL-4, IL-10, IL-13) and pro-inflammatory markers (IL-1β, IL-5, IFNγ, TNFα, IL-6, and KC/GRO), indicating it represents a broad inflammatory signature. PC2 explained an additional 12.7% of the variance and highlighted distinct cytokine dynamics, particularly strong associations with KC/GRO and IL-6, and inverse relationships with TNFα and IL-13. There were trends for Parity x Diet x Treatment interaction: F_(1,51)_ = 3.648, *p* = 0.062, □p^2^ = 0.067 for PC1). Genotype x Diet x Parity interaction: (F_(1,51)_ = 3.018, *p* = 0.088, □p^2^ = 0.056; **Figure 5E** & **F**). But no differences between groups were found in PC2 (*p* ≥ 0.146; **Figure 5G** & **H**).

Collectively, these findings indicate that E2 treatment significantly increases hippocampal inflammatory load in parous hAPOE□4 females in the dorsal hippocampus, particularly under a WD, as reflected by elevated PC1 scores representing a broad pro- and anti-inflammatory cytokine signature; in the ventral hippocampus although similar trends emerged, they were not significant.

### 2.4 PSD-95 expression did not differ by genotype, parity, E2 treatment

To assess whether genotype, parity, diet, or E2 treatment influence PSD-95, a synaptic protein, we examined levels of PSD-95 in both the dorsal and ventral hippocampus. In the dorsal hippocampus, there was a significant interaction between Genotype, Parity and Treatment (Genotype x Parity x Treatment: F_(1,65)_ = 4.983, *p* = 0.029, □p^2^ = 0.071; **Supplementary Figure 3A**), although no significant post hoc comparisons were found (*p* ≥ 0.203). Furthermore, there were no other significant main effects of interactions (*p* ≥ 0.129). We found no significant main effects or interactions in the ventral hippocampus (*p* ≥ 0.275; **Supplementary Figure 3B**).

### 2.5 Previous parity, but not E2 treatment, increased DCX-expression in WT rats whereas a WD decreased DCX expression based on genotype- and previous parity

We investigated the density of DCX-expressing cells in the dorsal and ventral dentate gyrus to assess neurogenesis. In the dorsal region, previous parity was associated with increased neurogenesis in WT rats fed a SD (E2 treated p=0.019) but decreased neurogenesis in hAPOE□4 fed a WD (*p*=0.003, Cohen’s *d* = 0.981). E2 treatment did not significantly influence neurogenesis in the WT SD-fed rats. However, in the hAPOE□4 rats, E2 decreased the density of DCX-expressing cells in nulliparous rats under a SD (*p*=0.002, Cohen’s *d* =1.015; Parity x Diet x Genotype x Treatment: F_(1, 61)_ = 5.00, *p* = 0.029, □p² = 0.076). A WD influenced neurogenesis but only in certain E2-treated rats, with WD exposure decreasing neurogenesis in WT primiparous rats (*p*=0.006, Cohen’s *d* = 1.292) yet increasing neurogenesis in hAPOE□4 nulliparous rats (*p*=0.0003 Cohen’s *d* = 1.448), compared to their SD counterparts. There were also significant main effect of genotype, as hAPOE□4 females had higher DCX levels than WT females (Genotype: F_(1, 61)_ = 6.80, *p* = 0.011, □p² = 0.078 ; **Figure 6C**) and significant interactions (Parity x Diet: F_(1, 61)_ = 7.88, *p* = 0.007, □p² = 0.011; Genotype x Diet x Treatment: F_(1, 61)_ = 10.51, *p* = 0.0019, □p² = 0.015; Parity x Diet x Treatment: F_(1, 61)_ = 10.400, *p* = 0.002, □p² = 0.015). However, no other significant main effects or interactions effects achieved significance (p’s > 0.18).

In the ventral dentate gyrus, as expected, parity increased neurogenesis under SD fed conditions in oil-treated WT females (*p* = 0.038, one-tailed, Cohen’s *d* = 0.820) with no significant effects with WD (*p* = 0.46, Cohen’s *d* = 0.773). E2 treatment increased neurogenesis in SD fed primiparous hAPOE□4 rats (p=0.004, Cohen’s *d* =1.059) compared to their oil control counterparts (Parity x Diet x Genotype x Treatment: F_(1, 55)_ = 10.48, *p* = 0.002, □p² = 0.16 **Figure 6D**), which was not observed under a WD. E2 decreased neurogenesis in primiparous hAPOE□4 rats compared to oil (p=0.013, not significant after Bonferroni correction). E2 did not significantly affect DCX-expressing cells in the WT animals under any condition. A WD decreased ventral neurogenesis in E2 treated primiparous hAPOE□4 rats only (*p* < 0.0001, Cohen’s *d* =1.648). There were also significant interactions (Parity x Diet: F_(1, 55)_ = 4.50, *p* = 0.038, □p² = 0.076; Parity x Diet x Treatment: F_(1, 55)_ = 5.01, *p* = 0.029, □p² = 0.083; Genotype x Diet x Treatment: F_(1, 55)_ = 4.73, *p* = 0.034, □p² = 0.079). No other significant main effects interactions achieved significance (p’s > 0.06).

## 3. Discussion

Our findings indicate that a WD in middle age increased body weight, certain metabolic hormones, and reduced neuroplasticity, while body weight influenced contextual memory. The ability of a WD or body weight to alter these phenotypes, however, depended on previous parity and APOE genotype. E2 treatment in middle age reduced WD-induced body weight gain and reduced WD-induced increases in metabolic hormones in WT rats but these effects were blunted or reversed in primiparous hAPOE□4 females, underscoring the influence of both genetic and reproductive factors on E2’s efficacy. Indeed, previous parity increased vulnerability to diet- and hormone-induced cognitive impairments, hippocampal inflammatory signaling and reduced neurogenesis, particularly in hAPOE□4 carriers. Importantly, previous parity, when combined with metabolic stress and genetic risk, may shift E2’s effects from neuroprotective to detrimental, including reduced support for hippocampal plasticity.

Specifically, WD consumption led to increases in circulating insulin, C-peptide, glucagon, and leptin alongside reduced PYY and GLP-1 levels. Notably, the metabolic effects of E2 treatment were genotype dependent as E2 reduced plasma glucagon in WT females but had no effect in hAPOE□4 females indicating that genotype, in part, shapes E2’s capacity to modulate metabolic hormones. At the cognitive level, body weight differentially predicted hippocampal-dependent memory depending on genotype and reproductive history. Increased body weight was associated with enhanced contextual fear memory in primiparous WT females but was associated with impaired contextual memory in primiparous hAPOE□4 females, a pattern that suggests genotype-dependent effects of metabolic burden on cognitive outcomes. Furthermore, hAPOE□4 females demonstrated superior cued fear conditioning on a standard diet, yet this was completely abolished following WD, indicating heightened sensitivity of hAPOE□4 females to metabolic stress. The most striking findings emerged in primiparous hAPOE□4 females, where WD and E2 treatment converged to amplify certain pathological outcomes. Under a WD, primiparous hAPOE□4 females exhibited elevated hippocampal inflammatory cytokine signatures in the dorsal hippocampus and reduced neurogenesis in the dorsal dentate gyrus. Collectively, these results demonstrate that the interplay between metabolic stress, genetic susceptibility, and reproductive history shapes brain aging trajectories in middle-aged females, with the primiparous hAPOE□4 phenotype representing a particularly vulnerable state.

### 3.1. Parity modulates susceptibility to WD-induced weight gain and metabolic hormone secretion whereas genotype affects the impact of E2 on metabolic hormone levels

Not surprisingly, WD consumption during middle age led to weight gain regardless of genotype or previous parity. Although parity is associated with increased weight gain in postmenopausal women (98), the number of childbirths plays an important role in obesity risk. Specifically, human females with one or two childbirths tend to have lower mean body weight and a reduced risk of obesity, whereas those with three or more childbirths exhibit higher body weight and increased obesity risk compared to nulliparous females (99,100). In the present study, we only examined primiparous rats. Importantly, E2 treatment mitigated WD-induced weight-gain in WT females, consistent with previous findings (101–103), however the most pronounced effect was E2’s ability to reduce weight gain in primiparous WT rats. In the present study, there were no beneficial effects of E2 to reduce body weight under a WD in hAPOE□4 females.

In addition to weight gain, WD consumption increased the levels of insulin, leptin, and C-peptide in plasma, indicating metabolic dysregulation as demonstrated previously (104). Furthermore, levels of the appetite-suppressing hormones PYY and GLP-1 were reduced in WD-fed females, which may contribute to overconsumption, increased body weight, and impaired glucose homeostasis (71,105–109). In addition, the principal component analyses indicated that E2 treatment altered metabolic hormones primarily in the WT females, suggesting that E2 was less effective in the hAPOEe4 females to reduce metabolic hormones. Together, these data indicate that previous parity and genotype affect the potency of E2 to improve metabolic regulation with respect to certain metabolic hormones and weight gain. In addition, previous parity may result in long-term metabolic changes which serve to buffer hormone responses to metabolic challenges later in life, particularly in WT animals.

### 3.2. Higher body weight and WD-diet exposure reduced contextual fear memory performance in primiparous hAPOE□4 females

Higher body weight was also associated with reduced associative memory in primiparous hAPOEε4 females, whereas the opposite was found in primiparous WT females. Previous reproductive experience is associated with improved hippocampus-dependent learning in middle aged WT rodents (110,111), and increased non-spatial strategy use in previous parous hAPOEe4 middle-age female rats(112), our finding are largely consistent with human studies that have shown improvements in cognition in middle to older age in previously parous females (113,114).

However, this relationship can change with dementia diagnosis or risk for dementia, as APOEe4 genotype and number of children was negatively associated with reaction time scores later in life (113) and the number of children had was negatively associated with verbal memory scores in late life but only in those diagnosed with dementia and not in those with normal cognitive aging or amnestic mild cognitive impairment (112). Together, these suggest that parity and genotype may interact to influence multiple domains of cognitive performance. Our findings further demonstrate that body weight modulated hippocampus-dependent learning in middle-aged females, emphasizing the interplay between reproductive experience, genotype, and metabolic status in shaping cognitive aging.

In this study, we found that SD hAPOE□4 females performed better in the amygdala-based associative memory task at middle age compared to WT. Previous studies have similarly reported better cognitive performance in younger adults, or no difference in middle age, in cognitively normal populations with APOEe4 genotype (115,116), indicating that compensatory mechanisms may be present at this age. This genotype difference, however, was only present in SD-fed subjects, as hAPOE□4 females exposed to WD presented with similar levels as WT WD-females. Emerging evidence from rodent models suggests that high-fat diets, and resulting obesity, impact spatial memory, especially in hAPOEε4 carriers, potentially by increasing central inflammation, promoting glucose intolerance, and Aβ pathology (117–120). Our findings that a WD may be detrimental to associative memory in APOE□4, is consistent with other studies in mice (121). One study reported that female mice benefit cognitively from dietary weight-loss interventions, suggesting dietary importance on cognition in females (122). Furthermore, another study demonstrated that high-fat diet exposure at middle age impaired spatial learning and memory in hAPOE□4, compared to WT, female mice, partially consistent with our findings (123). In human females, adherence to a Mediterranean diet was associated with reduce AD endophenotypes particularly in APOEe4 carriers, partially consistent with our findings (124). Taken together, in addition to their independent effects, obesity, parity and APOE genotype appear to interact to influence associative memory, suggesting that hAPOEε4 carriers may be more sensitive to diets high in sugar and/or fat, or increased body weight, an interaction that could further contribute to AD risk.

### 3.3. E2-induced central inflammation in hAPOE□4 primiparous females, especially under WD conditions

In hAPOE□4 rats, we found that primiparity was associated with decreased inflammatory load in the dorsal hippocampus, compared to nulliparity, an effect that was reversed in WT females, This is also partially consistent with another study that showed that previous parity in middle-aged hAPOEε4 genotype resulted in heightened inflammatory Th1/Th2 ratio in the hippocampus of middle-aged female rats (67). In individual cytokines, IL-1β, IFN-γ and IL-5, showed decreased expression in in primiparous compared to nulliparous rats in the ventral hippocampus (Supplementary Figure 1). This is in line with studies indicating a protective effect of previous parity in the hippocampus (51,53–56,58,60–63,125). In past studies, previous parity alters the hippocampal-inflammatory system and peripheral cytokine signaling with age, which may be related to the amount of parity (61,62,126,127).

We also found an increased inflammatory load (IL-1β, IFN-γ) in the ventral hippocampus in WD-fed animals compared to those fed a SD, however, this was only present in WT primiparous females. This is in line with previous studies demonstrating that diets rich in fat and/or sugar can increase hippocampal inflammation, although parity was not analyzed in these previous studies (75,77,80,128). This data therefore further demonstrates that parity may influence hippocampal sensitivity to diet-induced inflammation.

We found increased inflammatory load in the dorsal hippocampus, in response to E2 treatment, in primiparous hAPOE□4 females fed a WD, compared to all other groups. This is somewhat consistent with past findings as E2 reduced the levels of anti-inflammatory cytokines in the dorsal hippocampus depending on the age of first pregnancy in middle age indicating that previous parity may alter the inflammatory response to E2 (129).

Taken together, E2 treatment increased inflammatory load in WD-fed, primiparous hAPOE□4 females, suggesting that under conditions of metabolic and reproductive stress, estradiol may exacerbate, rather than reduce, neuroinflammation, characteristic of the healthy cell bias (112). In contrast, primiparity alone was associated with reduced inflammation, but this protective effect was absent in hAPOE□4 carriers. These findings are consistent with the healthy cell bias of estrogen action, which proposes that estradiol is beneficial only when cellular environments are healthy. In a compromised context, such as with WD exposure and genetic risk, however, its actions may shift toward being detrimental (130).

### 3.4. Parity, APOEε4, diet, and E2 interact to shape hippocampal neurogenesis in middle age

Our findings indicate that adult hippocampal neurogenesis in middle-aged females is shaped by the combined influence of parity, APOE genotype, diet, and E2. In the dorsal dentate gyrus, females carrying hAPOEε4 showed higher levels of neurogenesis than WT consistent with previous studies (131,132). However, this effect was attenuated when animals were exposed to a WD, particularly in primiparous females. Furthermore, previous parity increased neurogenesis in dorsal and ventral hippocampus, but only in the WT rats fed a SD, as the effect was attenuated with a WD. This aligns with recent reports that parity induces long-lasting modifications in hippocampal plasticity, which can interact with later-life metabolic and hormonal status to influence cognitive trajectories (133). Furthermore, studies in female rats have demonstrated that APOEε4 carriers show altered network-level activation and connectivity patterns that are sensitive to reproductive history (134), supporting the notion that neurogenesis-related processes may be differentially regulated depending on both genotype and parity. Furthermore, another study demonstrated that E2’s effects on hippocampal plasticity and inflammation are contingent on reproductive experience, with E2 enhancing neuroplastic processes in some contexts while exacerbating inflammatory responses in others (134), which is in line with our results.

Collectively, these findings suggest that neurogenesis in middle age is modulated by the interaction of genetic risk, reproductive history, and metabolic state. All of which are factors that may determine hippocampal resilience or vulnerability to cognitive decline.

Previous studies have demonstrated increased PSD-95 levels after E2 treatment, indicating that E2 may enhance synaptic protein expression and plasticity (135–139); although another study found no effect (140). The literature presents mixed findings regarding the influence of the hAPOE□4 genotype on PSD-95 expression (141–144). It has been suggested that these differences may be due to compensatory mechanisms early in disease progression which fail as the disease progresses, leading to a subsequent decline in PSD-95 levels. Further studies with more experimental subjects are needed to further investigate the relationship between diet, parity and E2 treatment.

### 3.5 Limitations and Future Directions

The current study examined only primiparous females, which does not capture the full spectrum of reproductive experience. Prior research suggests that higher levels of parity, such as biparity or multiparity, may have differential or even detrimental effects on brain aging and AD risk (51,55–57). Including multiple levels of parity in future work may help clarify the nuanced ways reproductive history shapes neurobiological aging and interacts with genetic risk.

Future studies should also incorporate measures of key physiological mechanisms that may mediate these effects, including peripheral inflammatory and insulin sensitivity, brain insulin signaling, blood-brain barrier integrity, and pathological markers on AD, including tau and Amyloid-β, across parity and genotype could help to disentangle the pathways by which metabolic and hormonal factors influence brain health. A better understanding of these mechanisms may inform more personalized interventions for females at elevated genetic risk for AD with different reproductive experience.

### 3.6. Conclusions

This study provides novel insight into the complex interplay between reproductive history, APOE genotype, diet, and E2 treatment in shaping metabolic, cognitive, inflammatory, and neurogenic outcomes in middle-aged females. Our findings indicate that previous parity coupled with E2 treatment confers greater protection against weight gain and metabolic disruption under a WD but may also increase susceptibility to diet- and hormone-induced cognitive impairments, inflammation and reduced neurogenesis in hAPOE□4 carriers. Although E2 treatment showed some beneficial effects on metabolic markers, these effects were blunted or reversed in primiparous hAPOE□4 females, underscoring the influence of both genetic and reproductive factors on E2’s efficacy. Importantly, previous parity, when combined with metabolic stress and genetic risk, may shift E2’s effects from neuroprotective to detrimental. These findings emphasize the need for more personalized approaches to hormone therapy and brain health in females, considering genotype, reproductive history, and lifestyle factors such as diet.

## Acknowledgements

We would like to thank Peixin Shi, Aya Chahrour, Ziyi Zhang, Valerie Cheng, and Lele Ma for their valuable assistance with scoring behavioral videos. We are also grateful to Bonnie Lee for generously sharing her expertise and training in these analytical methods. Thank you also to Melike Cevizci for assisting with hormone injections. Their contributions were instrumental to the completion of this work.

## Author contributions

**JER**: Conceptualization; Methodology; Investigation; Formal analysis; Data curation; Visualization; Writing – original draft. **AM:** Investigation, Formal analysis; Writing. **SEL, KAG, SW**: Investigation. **RKR** & **TFLS**: Investigation; Data curation; Supervision. **SB**-Conceptualization; Funding acquisition, Writing-review and editing; **LAMG**: Conceptualization; Funding acquisition; Supervision; Writing – review & editing; Project administration.

## Data availability

Data will be made available upon reasonable request.

## Supplementary

**Supplementary Figure 1.**
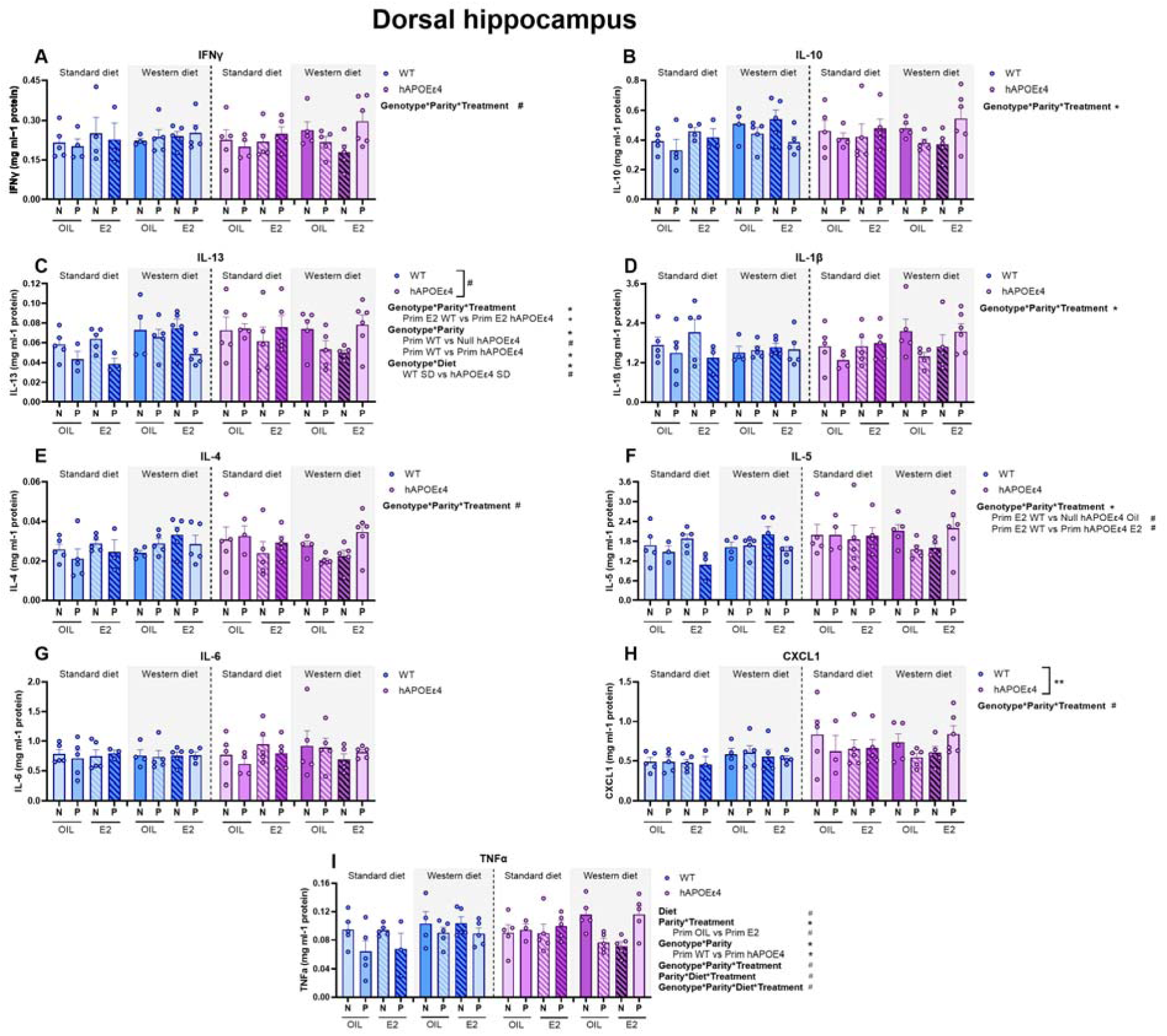
Cytokines in the dorsal hippocampus. Levels of interferon γ (IFNγ; A), interleukin (IL)-10 (B), IL-13 (C), IL-1β (D), IL-4 (E), IL-5 (F), IL-6 (G), chemokine (C-X-C motif) ligand 1 (CXCL1; H), tumor necrosis factor (TNF) α (I). n = 3-6 for treatment, genotype, parity and diet groups. *\** p *< 0.05, \*\** p *<* 0.01, # p < 0.10. N = nulliparous; P = primiparous; WT = wildtype; E2 = estradiol.

**Supplementary Figure 2.**
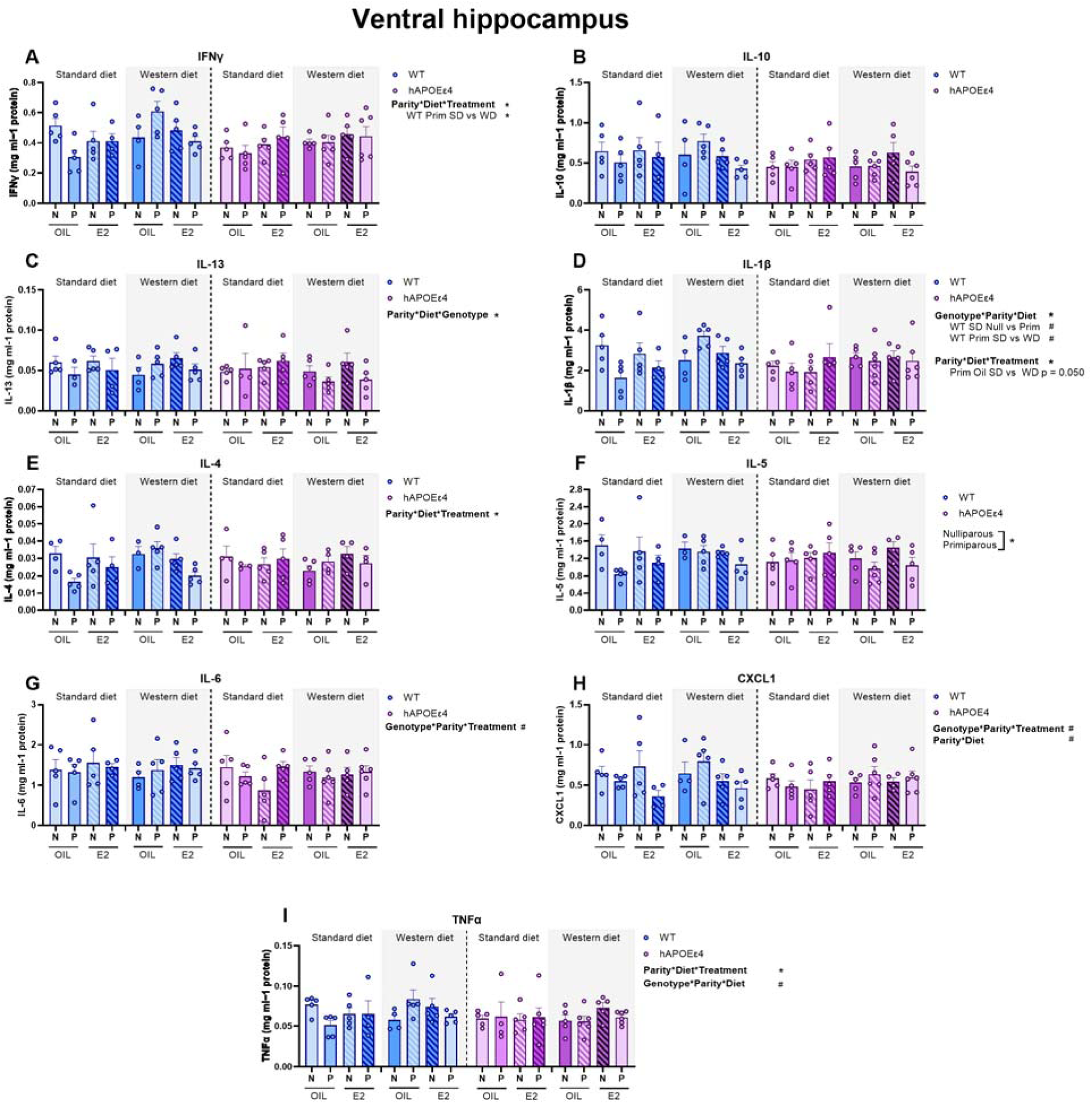
Cytokines in the ventral hippocampus. Levels of interferon γ (IFNγ; A), interleukin (IL)-10 (B), IL-13 (C), IL-1β (D), IL-4 (E), IL-5 (F), IL-6 (G), chemokine (C-X-C motif) ligand 1 (CXCL1; H), tumor necrosis factor (TNF) α (I). n = 3-6 for treatment, genotype, parity and diet groups. *\** p *< 0.05,* # p < 0.10. N = nulliparous; P = primiparous; WT = wildtype; E2 = estradiol.

**Supplementary Figure 3.**
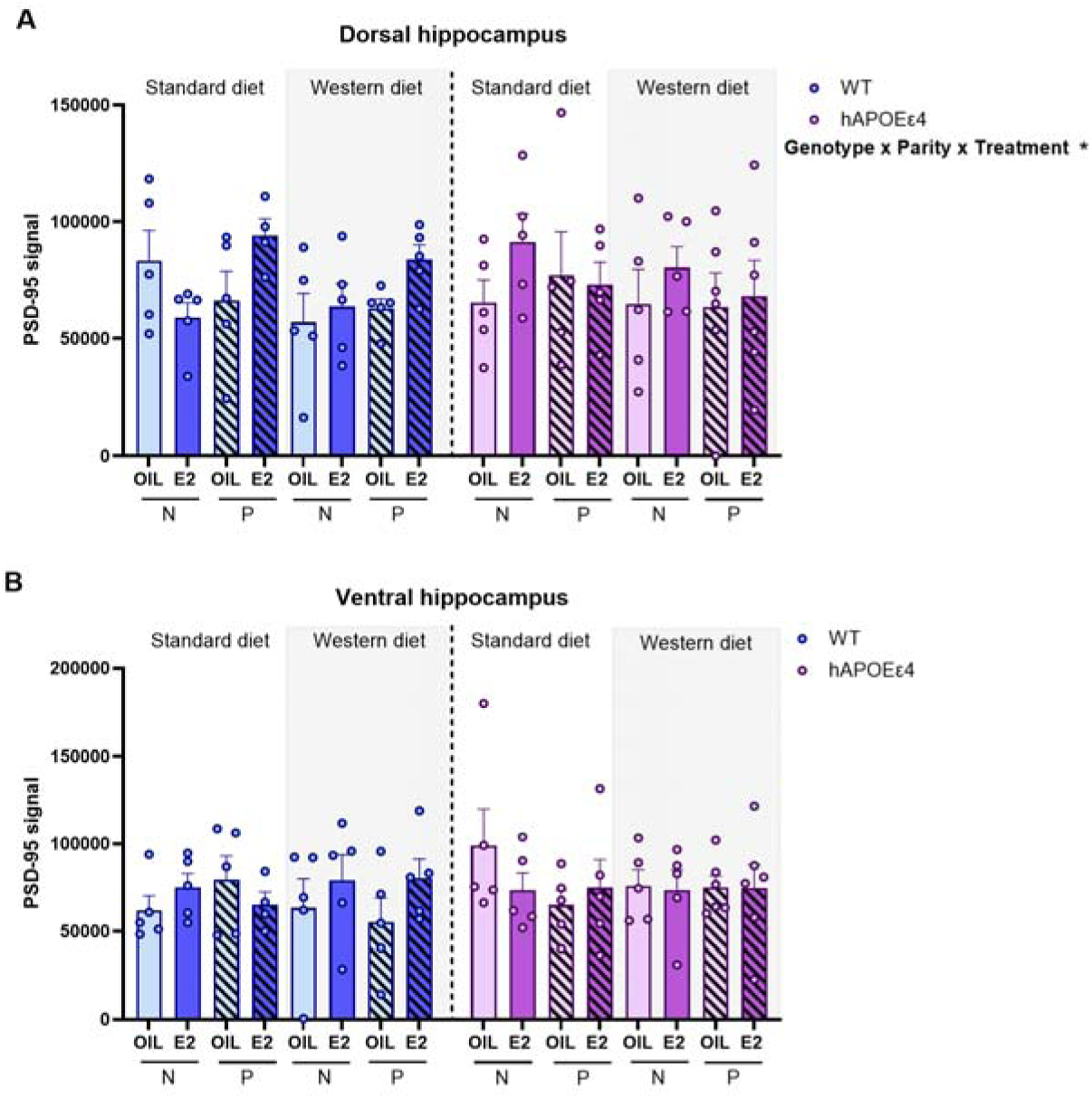
Effects of genotype, parity, diet, and E2 treatment on synaptic plasticity. PSD-95 signal in the dorsal (dHPC; A) and ventral hippocampus (vHPC: B) in middle-aged females. n = 4-6 for treatment, genotype, parity and diet groups. * p < 0.05, ** p < 0.005 # < 0.100. E2 = estradiol, WT = wildtype, SD= standard diet, WD = western diet, N = nulliparous, P = primiparous.

**Supplementary Figure 1.**
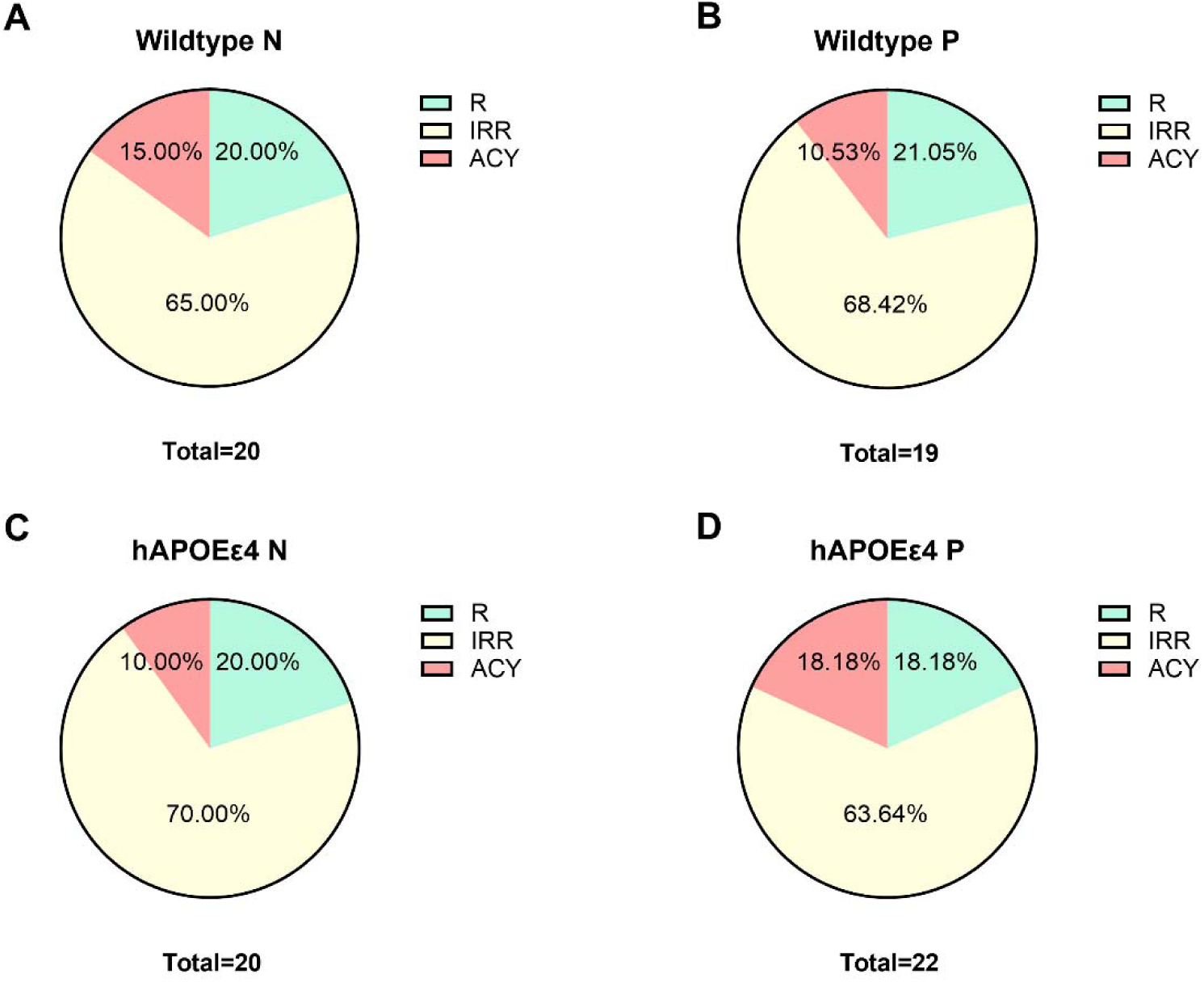
Estrous cycling before treatment (10-11 months of age). Cycling in wildtype (WT) nulliparous (N; A), and primiparous (P; B), and in hAPOE□4 N (C) and hAPOE□4 P (D).

**Supplementary Figure 2.**
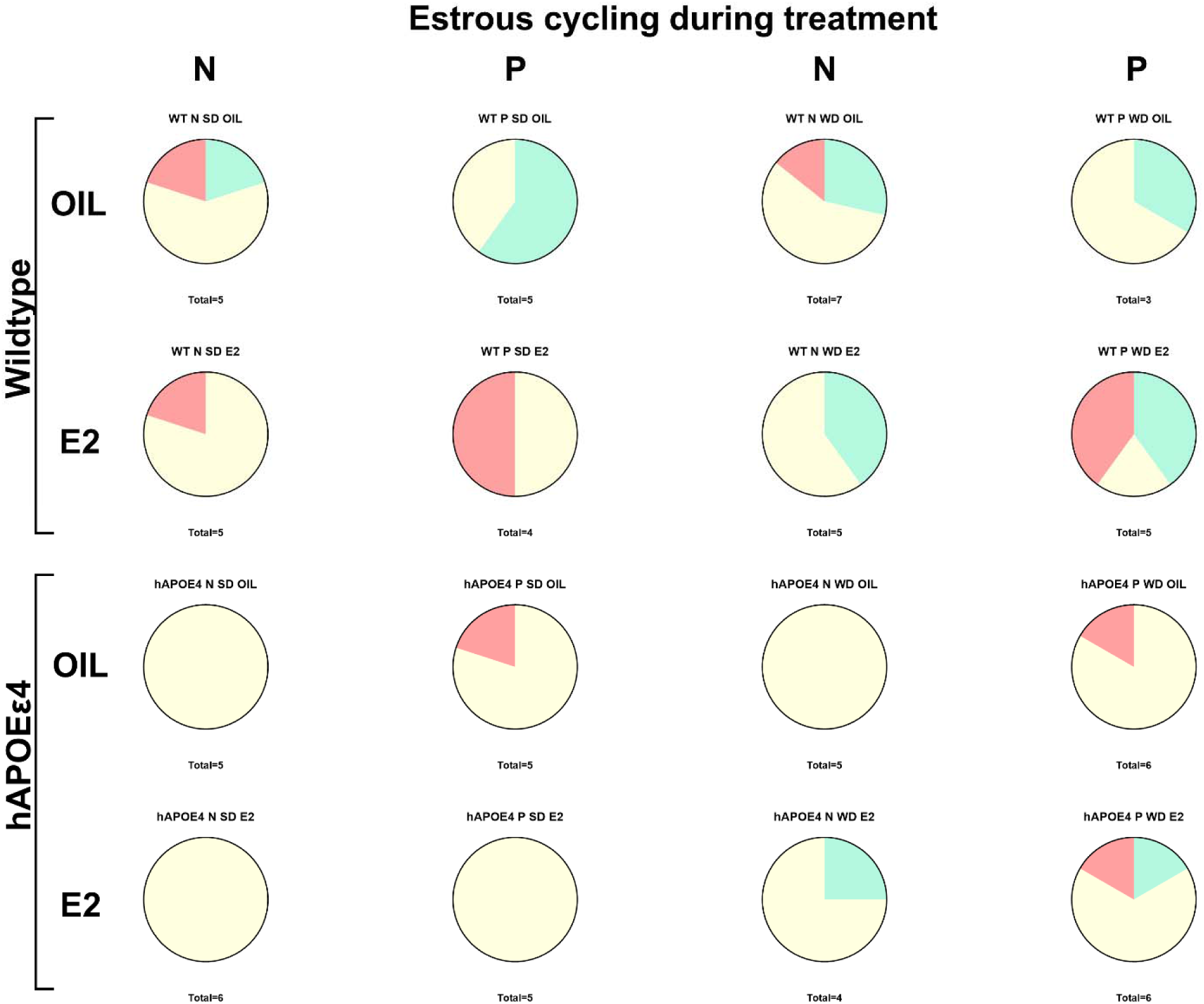
Estrous cycling during treatment (11-12 months of age). Cycling in wildtype (WT) and hAPOE□4 females of nulliparous (N) or primiparous (P) parity, fed with Standard (SD) or western diet (WD) and treated with estradiol (E2) or vehicle (oil).

**Supplementary Figure 3.**
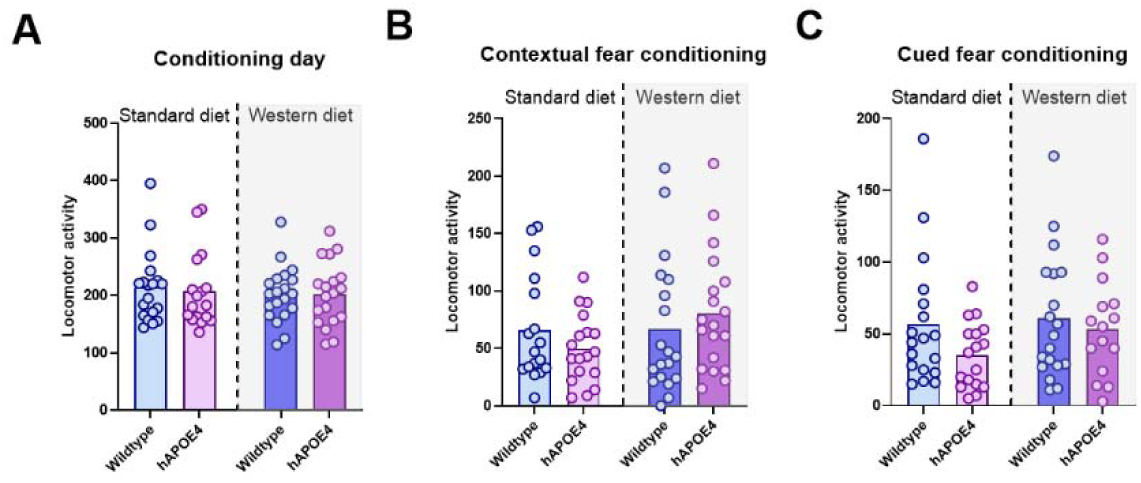
No significant differences in locomotor activity were found between genotypes or diet groups during the fear conditioning task. Locomotor activity during the conditioning day (A), contextual (B) and cued (C) fear conditioning tasks. n = 17-20.

**Table 1.** Statistical details of cytokines loaded onto Principal Component (PC) 1 and 2 in the dorsal and ventral hippocampus.

| PC1 | r | p |
| --- | --- | --- |
| IL-4 | 0.89 | 4.01×10<sup>-25</sup> |
| IL-10 | 0.85 | 1.13×10<sup>-20</sup> |
| IL-5 | 0.83 | 2.67×10<sup>-19</sup> |
| IL-13 | 0.82 | 4.05×10<sup>-18</sup> |
| IFNγ | 0.77 | 2.76×10<sup>-15</sup> |
| IL-1β | 0.72 | 1.88×10<sup>-12</sup> |
| TNFα | 0.65 | 7.03×10<sup>-10</sup> |
| CXCL1 | 0.65 | 1.26×10<sup>-09</sup> |
| IL-6 | 0.43 | 1.75×10<sup>-04</sup> |

| PC2 | r | p |
| --- | --- | --- |
| IL-6 | 0.75 | 6.00×10<sup>-14</sup> |
| CXCL1 | 0.39 | 8.70×10<sup>-04</sup> |
| IFNγ | 0.28 | 1.89×10<sup>-02</sup> |
| IL-1β | 0.25 | 3.60×10<sup>-02</sup> |
| IL-5 | -0.34 | 3.73×10<sup>-03</sup> |
| IL-13 | -0.35 | 2.84×10<sup>-03</sup> |

Ventral hippocampus
| PC1 | r | p |
| --- | --- | --- |
| IL-4 | 0.89 | 1.10×10<sup>-23</sup> |
| IL-1β | 0.88 | 1.58×10<sup>-22</sup> |
| IL-5 | 0.87 | 1.53×10<sup>-21</sup> |
| IFNγ | 0.85 | 4.87×10<sup>-20</sup> |
| IL-10 | 0.82 | 1.58×10<sup>-17</sup> |
| TNFα | 0.72 | 5.93×10<sup>-12</sup> |
| IL-13 | 0.69 | 1.48×10<sup>-10</sup> |
| CXCL1 | 0.53 | 3.59×10<sup>-06</sup> |
| IL-6 | 0.34 | 5.00×10<sup>-03</sup> |

| PC2 | r | p |
| --- | --- | --- |
| IL-6 | 0.68 | 3.25×10<sup>-10</sup> |
| CXCL1 | 0.55 | 1.75×10<sup>-06</sup> |
| IFNγ | -0.37 | 2.04×10<sup>-03</sup> |
| IL-1β | -0.47 | 5.27×10<sup>-05</sup> |

