## Supplement for "Reproductive history differentially shapes the neural response of middle-aged females to estradiol therapy after a metabolic challenge"

**Supplementary**

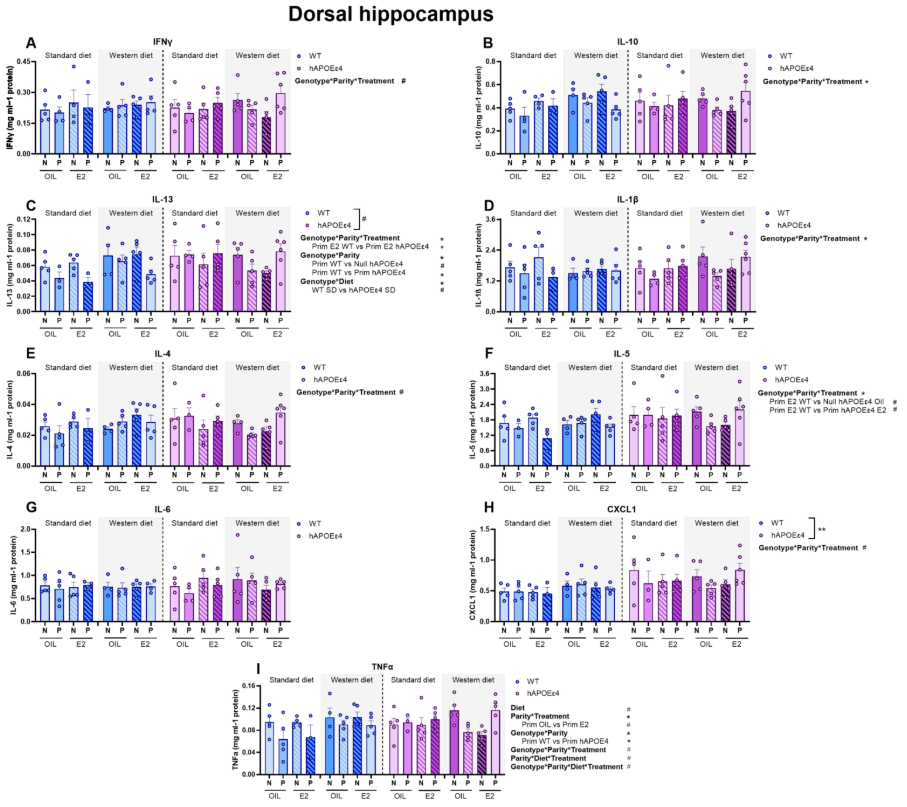

*Supplementary Figure 1.* **Cytokines in the dorsal hippocampus.** Levels of interferon γ (IFNγ; A), interleukin (IL)-10 (B), IL-13 (C), IL-1β (D), IL-4 (E), IL-5 (F), IL-6 (G), chemokine (C-X-C motif) ligand 1 (CXCL1; H), tumor necrosis factor (TNF) α (I). n = 3-6 for treatment, genotype, parity and diet groups. *** p *< 0.05, *** p *<* 0.01, # p < 0.10. N = nulliparous; P = primiparous; WT = wildtype; E2 = estradiol.

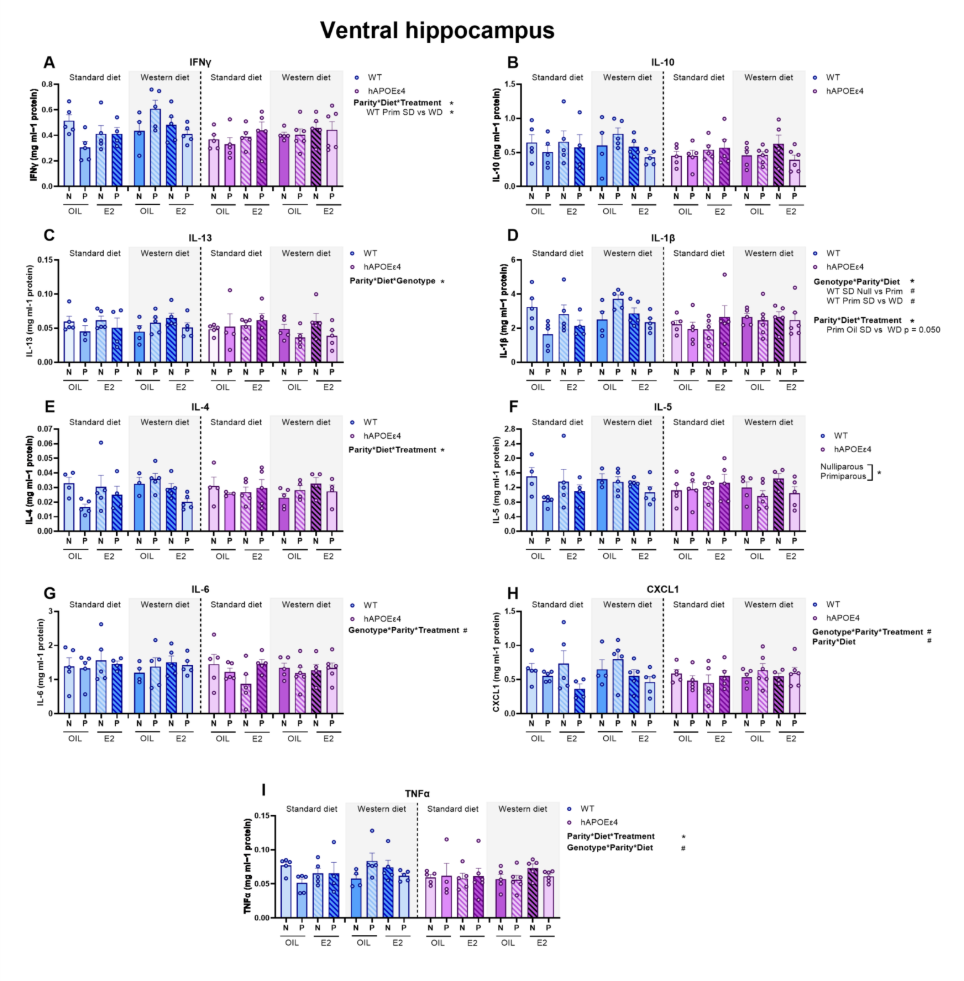

*Supplementary Figure 2.* **Cytokines in the ventral hippocampus.** Levels of interferon γ (IFNγ; A), interleukin (IL)-10 (B), IL-13 (C), IL-1β (D), IL-4 (E), IL-5 (F), IL-6 (G), chemokine (C-X-C motif) ligand 1 (CXCL1; H), tumor necrosis factor (TNF) α (I). n = 3-6 for treatment, genotype, parity and diet groups. *** p *< 0.05,* # p < 0.10. N = nulliparous; P = primiparous; WT = wildtype; E2 = estradiol.

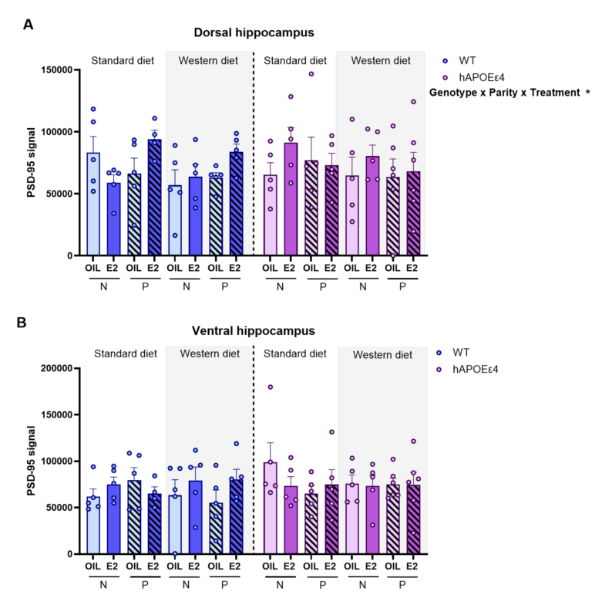

*Supplementary Figure 3.* **Effects of genotype, parity, diet, and E2 treatment on synaptic plasticity.** PSD-95 signal in the dorsal (dHPC; A) and ventral hippocampus (vHPC: B) in middle-aged females. n = 4-6 for treatment, genotype, parity and diet groups. * p < 0.05, ** p < 0.005 # < 0.100. E2 = estradiol, WT = wildtype, SD= standard diet, WD = western diet, N = nulliparous, P = primiparous.

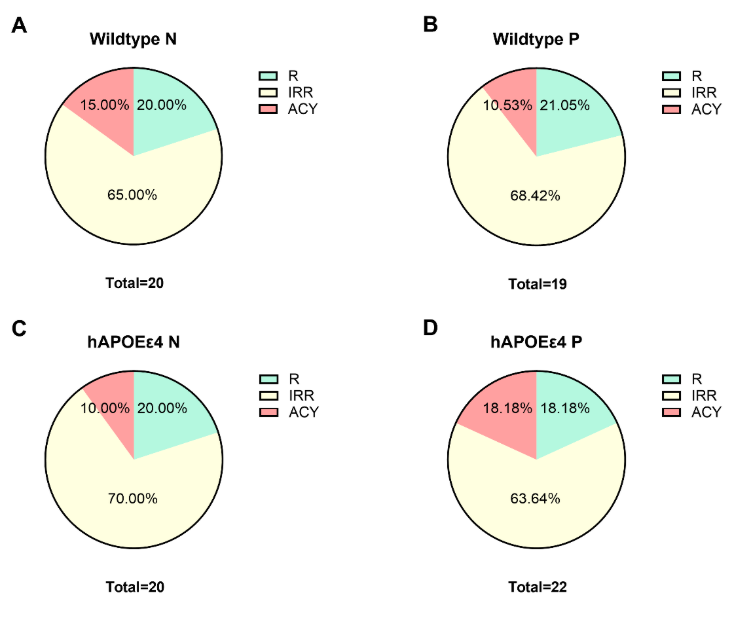

*Supplementary Figure 4.* **Estrous cycling before treatment (10-11 months of age).** Cycling in wildtype (WT) nulliparous (N; A), and primiparous (P; B), and in hAPOEɛ4 N (C) and hAPOEɛ4 P (D).

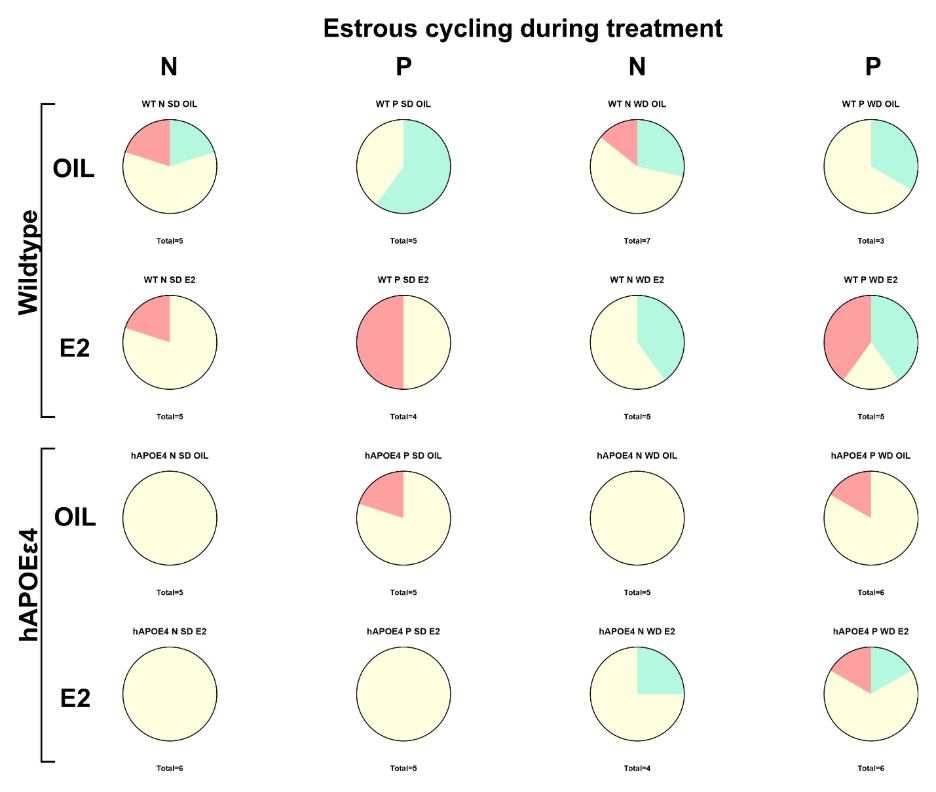

*Supplementary Figure 5.* **Estrous cycling during treatment (11-12 months of age).** Cycling in wildtype (WT) and hAPOEɛ4 females of nulliparous (N) or primiparous (P) parity, fed with Standard (SD) or western diet (WD) and treated with estradiol (E2) or vehicle (oil).

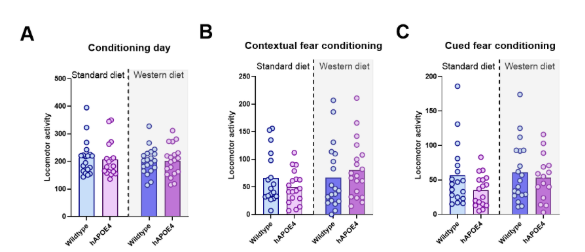

*Supplementary Figure 6.* **No significant differences in locomotor activity were found between genotypes or diet groups during the fear conditioning task.** Locomotor activity during the conditioning day (A), contextual (B) and cued (C) fear conditioning tasks. n = 17-20.

*Table 1* Statistical details of cytokines loaded onto Principal Component (PC) 1 and 2 in the dorsal and ventral hippocampus.

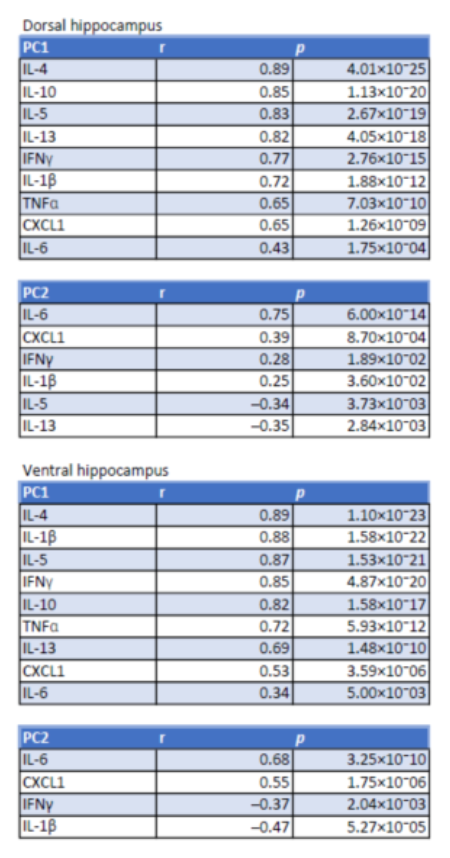
